# How does the brain navigate knowledge of social relations? Testing for shared neural mechanisms for shifting attention in space and social knowledge

**DOI:** 10.1101/2020.06.20.162255

**Authors:** Meng Du, Ruby Basyouni, Carolyn Parkinson

## Abstract

How does the human brain support reasoning about social relations (e.g., social status, friendships)? Converging theories suggest that navigating knowledge of social relations may co-opt neural circuitry with evolutionarily older functions (e.g., shifting attention in space). Here, we analyzed multivoxel response patterns of fMRI data to examine the neural mechanisms for shifting attention in knowledge of a social hierarchy. The “directions” in which participants mentally navigated social knowledge were encoded in multivoxel patterns in superior parietal cortex, which also encoded directions of attentional shifts in space. Exploratory analyses implicated additional regions of posterior parietal and occipital cortex in encoding analogous mental operations in space and social knowledge. However, cross-domain analyses suggested that attentional shifts in space and social knowledge may be encoded in functionally independent response patterns. These results elucidate the neural basis for navigating abstract knowledge of social relations, and its connection to more basic mental operations.

## Introduction

Effectively navigating the human social world requires tracking, encoding, and reasoning about the bonds, rivalries, and hierarchies that comprise it. Correspondingly, humans and other group-living primates possess sophisticated social cognitive abilities. One such ability involves learning and reasoning about others’ relative ranks in social hierarchies, which is important for strategically choosing allies and avoiding potentially harmful conflicts (e.g., Cheney and Seyfarth, 1990a). Rough, approximate assessments of someone’s power or status can be gleaned from perceptual cues (Hall et al., 2005; Marsh et al., 2009; Mattan et al., 2017), but more precise knowledge is achieved through observing and participating in encounters between pairs of individuals, then using transitive inference to ascertain the relative status of those whom one has not seen interacting with each other (Cheney and Seyfarth, 1990a, 1990b; Gazes et al., 2017; Grosenick et al., 2007; Kumaran et al., 2012; Paz-y-Miño et al., 2004; White and Gowan, 2013). Indeed, there is evidence that humans and other highly social animals acquire nuanced knowledge of others’ relative status in social hierarchies in this way (Cheney and Seyfarth, 1990a, 1990b; Kumaran et al., 2012), and use such knowledge strategically, such as when choosing social partners (Cheney and Seyfarth, 1990a). Past research has examined the neural basis of how people learn about social hierarchies and the neural correlates of social status (e.g., Kumaran et al., 2012; Muscatell et al., 2012; Zink et al., 2008). Yet, relatively little is known about the neural mechanisms that support reasoning about social relations, or how they relate to less abstract mental operations.

### Do Mental Representations of Space Scaffold Those of Social Relations?

Theories from neuroscience, psychology, and cognitive linguistics have highlighted widespread parallels in how people speak and think about space and social knowledge, and have inspired suggestions that mentally navigating social structures might have co-opted pre-existing neural architecture that originally supported evolutionarily older functions, such as representing and navigating physical space. For example, Conceptual Metaphor Theory suggests that the language used to describe abstract information can shed light on how the mind processes such information (Lakoff and Johnson, 1980). In particular, the Spatialization of Form Hypothesis (Lakoff, 1987) suggests that the widespread use of spatial language to describe conceptual relations, such as social relations, reflects the spatial organization of knowledge of such relations. Thus, phrases like “top of the pecking order” and “low status” may not just be figures of speech, but rather, figures of thought that illuminate the structure of underlying mental representations (Lakoff, 1986). Research findings and theoretical perspectives from cognitive neuroscience have converged with these ideas, suggesting that many cognitive capacities emerge through the repurposing of existing neural architecture, both over the course of evolution and in the case of culturally learned capacities, through development (Anderson, 2010; Dehaene and Cohen, 2007; Parkinson and Wheatley, 2015, 2013; Yamazaki et al., 2009). Such accounts provide potential neural mechanisms for theories from cognitive linguistics (e.g., Lakoff, 1987) and psychology (e.g., Williams et al., 2009) that suggest that representational resources with a precursory role in processing space were repurposed to process more abstract domains, such as time and social relations. For example, it has been suggested that, over the course of human evolution, regions of the posterior parietal cortex (PPC) originally devoted to representing and allocating attention in peripersonal space were co-opted to perform analogous operations on increasingly abstract contents (Yamazaki et al., 2009). Consistent with this possibility, findings from neuroimaging studies suggest considerable overlap in the brain regions that are recruited when processing spatial information and abstract social knowledge, particularly in the PPC (Parkinson and Wheatley, 2013; Yamazaki et al., 2009).

### Does Posterior Parietal Cortex Support Mentally “Navigating” Knowledge of Social Relations?

While past findings suggest that areas in the PPC play a role in processing knowledge of social relations, many questions remain. For example, a previous study found that regions of the inferior PPC were more active when people made more difficult social status comparisons (e.g., determining which of two naval officers had a higher rank, when their ranks were very similar) than for easier ones (e.g., determining which of two naval officers had a higher rank, when their ranks differed greatly; Chiao et al., 2009). Yet, it is difficult to ascertain if such effects reflect the neural representation of status knowledge or the more demanding nature of more difficult comparisons, especially given that the same brain region is also implicated in unrelated tasks when they demand attention or are difficult (Göbel et al., 2004; Shuman and Kanwisher, 2004). Thus, whereas the neural basis of *acquiring* knowledge about social relations (e.g., social status hierarchies) has been relatively well-characterized (Kumaran et al., 2012), further work is needed to understand how the brain supports reasoning about, or mentally “navigating” knowledge of social relations.

Addressing this question is likely to benefit from approaches that consider the information contained in spatial patterns of responses within brain areas, rather than analyses that only consider the overall magnitude of neural responses: Whereas many regions, including regions of the PPC, may be more active when tasks are more difficult or attention-demanding, regions that specifically encode the mental navigation of social knowledge should respond differently when mentally traversing knowledge of social relations in different ways, even when equating for difficulty. Multivoxel pattern analysis (MVPA) is likely to be especially fruitful in this context, given that it is sensitive to differences in the spatial patterning of responses in a brain region (e.g., for different kinds of stimuli or mental operations), even when the overall response magnitudes are equivalent (Peelen and Downing, 2007).

Here, to elucidate the neural basis of representing and mentally navigating social hierarchy knowledge (operationalized as shifting attention from one’s mental representation of one person to that of another person, based on one’s knowledge of social relations between people and task-based goals), we characterized the response patterns evoked when people reasoned about individuals’ relative location in a learned social hierarchy. Specifically, we used MVPA to compare neural response patterns on trials that were matched for difficulty, but that differed in the nature of mental navigation within participants’ knowledge of a set of social relations. The trials differed in the “directions” of navigation, such that participants shifted their attention towards either a more powerful person or a less powerful person in the social hierarchy. If response patterns within a brain region subserve mentally navigating internal representations of social hierarchies, then in that region, response patterns should be more similar for shifts of attention in the same “direction” than in different “directions” within knowledge of a social hierarchy.

Past research has highlighted the importance of the superior parietal lobule (SPL) in supporting shifts of attention both in the perceptual environment and in internal knowledge representations (Cabeza et al., 2008; Hutchinson et al., 2009; Knops et al., 2009; Serences et al., 2004). Thus, we focused on the SPL in our main analyses. In addition, given that relatively little is known about how the brain supports traversing knowledge of social relations, and given that past related research has tended to use mass univariate analyses, which are not sensitive to effects carried in multivoxel patterns, we also performed exploratory analyses across regions spanning all of cerebral cortex and subcortical gray matter structures.

### Are Shifts of Attention in Space and Knowledge of Social Relations Encoded in the Same Way?

As described above, considerable overlap has been documented in the brain regions that are recruited when processing knowledge of social hierarchies and other domains of content. Therefore, we also sought to test if in such regions, shifts of attention in knowledge of social relations (exemplified by status hierarchies here) are encoded similarly to shifts of attention in physical space. However, shared encoding mechanisms cannot be inferred from overlapping fMRI activations alone, as such results can be produced either from common codes for processing different domains or modalities of information (Parkinson et al., 2014; Peelen et al., 2010), or by spatially overlapping but functionally independent response patterns (Downing et al., 2007; Peelen et al., 2006). Thus, MVPA affords a more stringent test of whether overlapping fMRI activations reflect shared or distinct coding mechanisms (Peelen and Downing, 2007). For example, Knops et al. (Knops et al., 2009) trained a machine learning classifier only to discriminate fMRI response patterns in the SPL during overt leftward and rightward attentional shifts, and found that it correctly generalized to distinguishing brain responses during mental subtraction and addition. This suggests that mental arithmetic relies on neural mechanisms that also support shifting attention in space, and may involve mental operations akin to shifting attention along a horizontally oriented mental number line. Using MVPA in this manner can provide insight into the representational contents of brain regions, and assess if common encoding mechanisms support analogous mental operations in different domains of information.

### The Current Study

In the current study, we sought to elucidate the neural basis of shifting attention within knowledge of social relations (here, social hierarchy knowledge), and to test if common neural mechanisms are involved in shifting attention in space and internal representations of social relations. Participants first learned, through trial and error, the relative positions of 9 individuals in a fictive social hierarchy (Figs. 1 and 2). During scanning, participants performed a social hierarchy navigation task, where they determined who in the hierarchy was a given number of steps more or less powerful than a reference person (Fig. 3). Participants later performed an ostensibly unrelated eye movement task during fMRI scanning, which allowed us to characterize activity patterns associated with overt spatial shifts of attention. After scanning, participants performed an open-ended task probing the structure of their mental representations of the social hierarchy (Fig. 4). We tested if distributed response patterns in the SPL encode shifts of attention in knowledge of social relations. We also tested if shifts of attention in space and social relationship knowledge are encoded in overlapping brain regions, and if so, whether they are encoded in the same way in such regions. Testing these questions seeks to advance our understanding of how the human brain supports the ability to navigate knowledge of social relations, and more generally, how abstract feats of human social cognition relate to more concrete operations, such as processing the space around oneself.

**Figure 1.**
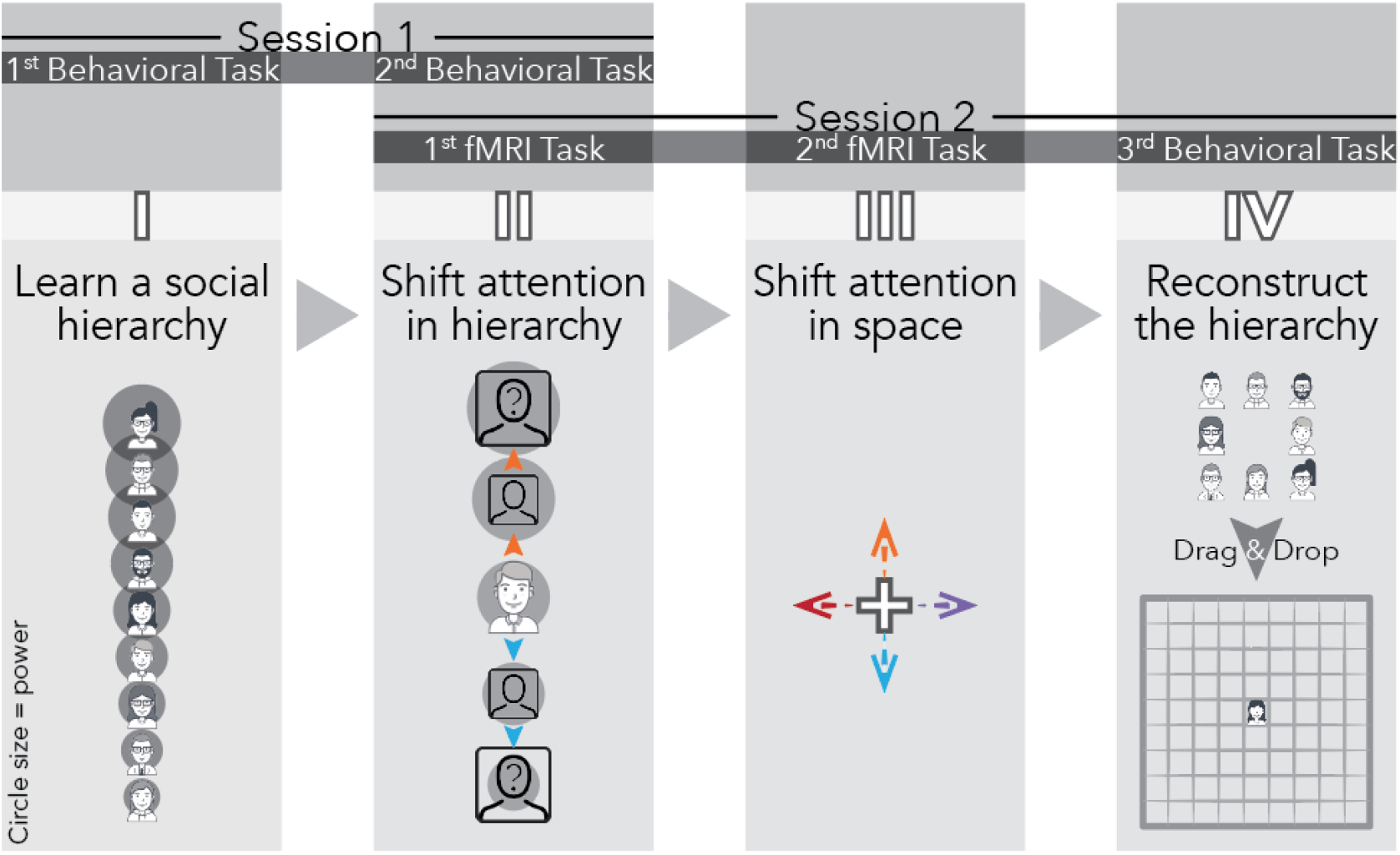
Overview of paradigm. Participants attended a behavioral session (Session 1) and an fMRI session (Session 2). In Session 1, they first learned a social hierarchy of 9 people through trial and error (Task I*). They then completed a social hierarchy navigation task (Task II*), where they were repeatedly asked to determine who was a given number of steps more or less powerful than another person in the learned social hierarchy. In Session 2, participants underwent fMRI scanning while performing the social task (Task II) again, followed by Task III, which involved overt shifts of attention in external space (i.e., eye movements). At the end of the experiment, we probed participants’ mental representations of the social hierarchy (i.e., mappings between positions in the hierarchy and locations in space) by asking them to arrange the faces in the way that they thought best reflected the people’s relative power in the organization (Task IV). * Note: in schematic illustrations of Tasks I and II, faces are depicted at different locations and within circles of varying sizes to indicate relative positions in the social hierarchy. These cues are included here only for illustration. This vertical arrangement, or any systematic spatial arrangement of the faces, was never shown to participants, nor were the faces depicted in circles of varying sizes. No spatial cues were provided to participants in Tasks I and II.

**Figure 2.**
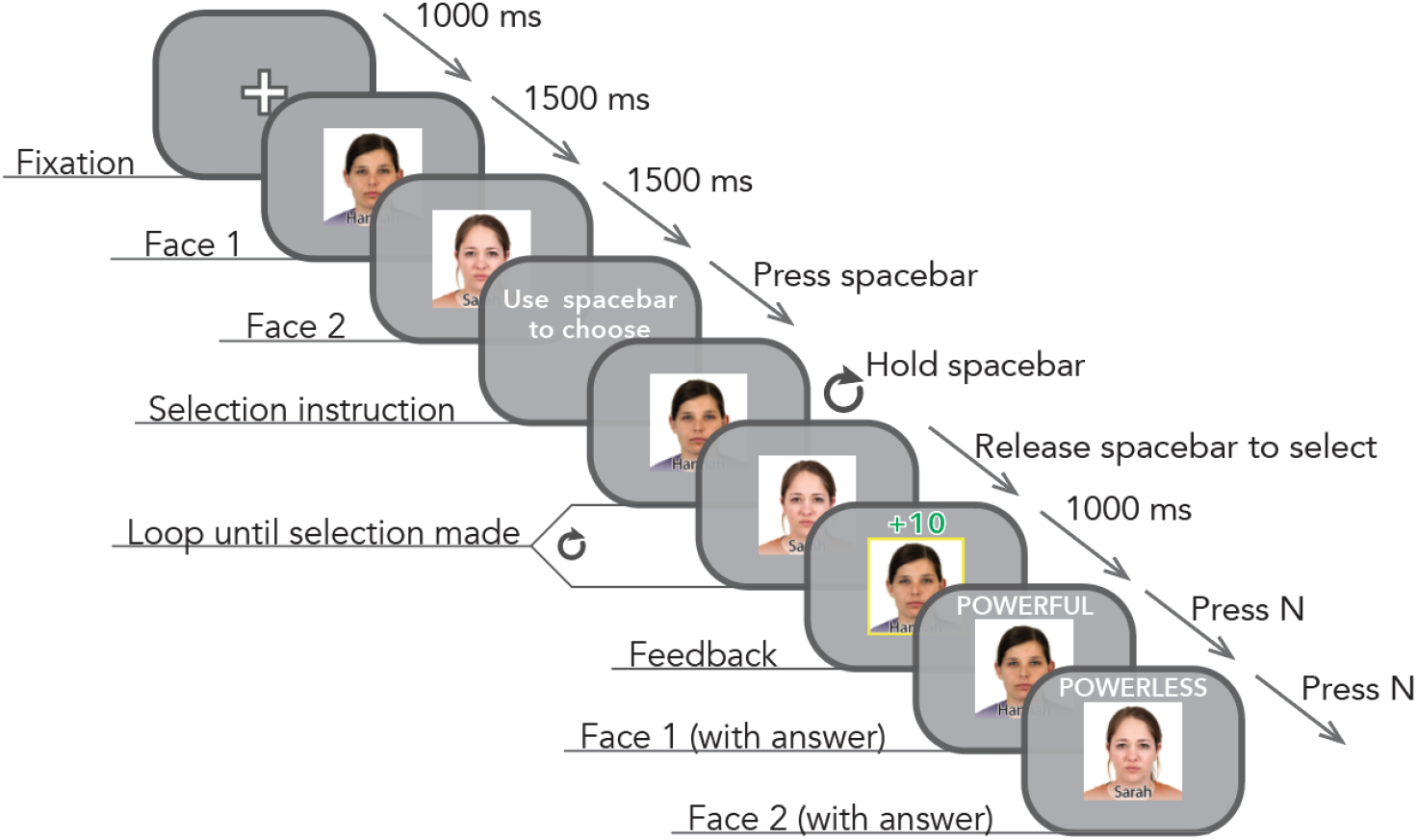
Learning the hierarchy (Task I). In this task, participants learned each person’s relative position in the social hierarchy through trial and error. To avoid biasing participants towards thinking of the hierarchy in spatial terms, or using any particular spatial mapping, both presentation of the faces and the response paradigm were designed to be sequential, rather than spatial (for example, faces and response keys were never arranged side-by-side or vertically). In each trial, two faces were presented on the screen one at a time, in a random order. Participants were then prompted to select the more powerful person in the pair by pressing and holding the space bar. While they held down the spacebar, the two face images were cycled through repeatedly in the same order as earlier in the trial. Participants chose one of the faces by releasing the space bar whenever they saw the person they intended to choose. The selected face was then highlighted together with a score (e.g., “+10 points” for correct responses, or “-10 points” for incorrect responses). Finally, the two faces were presented sequentially again, along with the correct information about the relative power of each face. Participants then pressed “N” to advance to the next trial.

**Figure 3.**
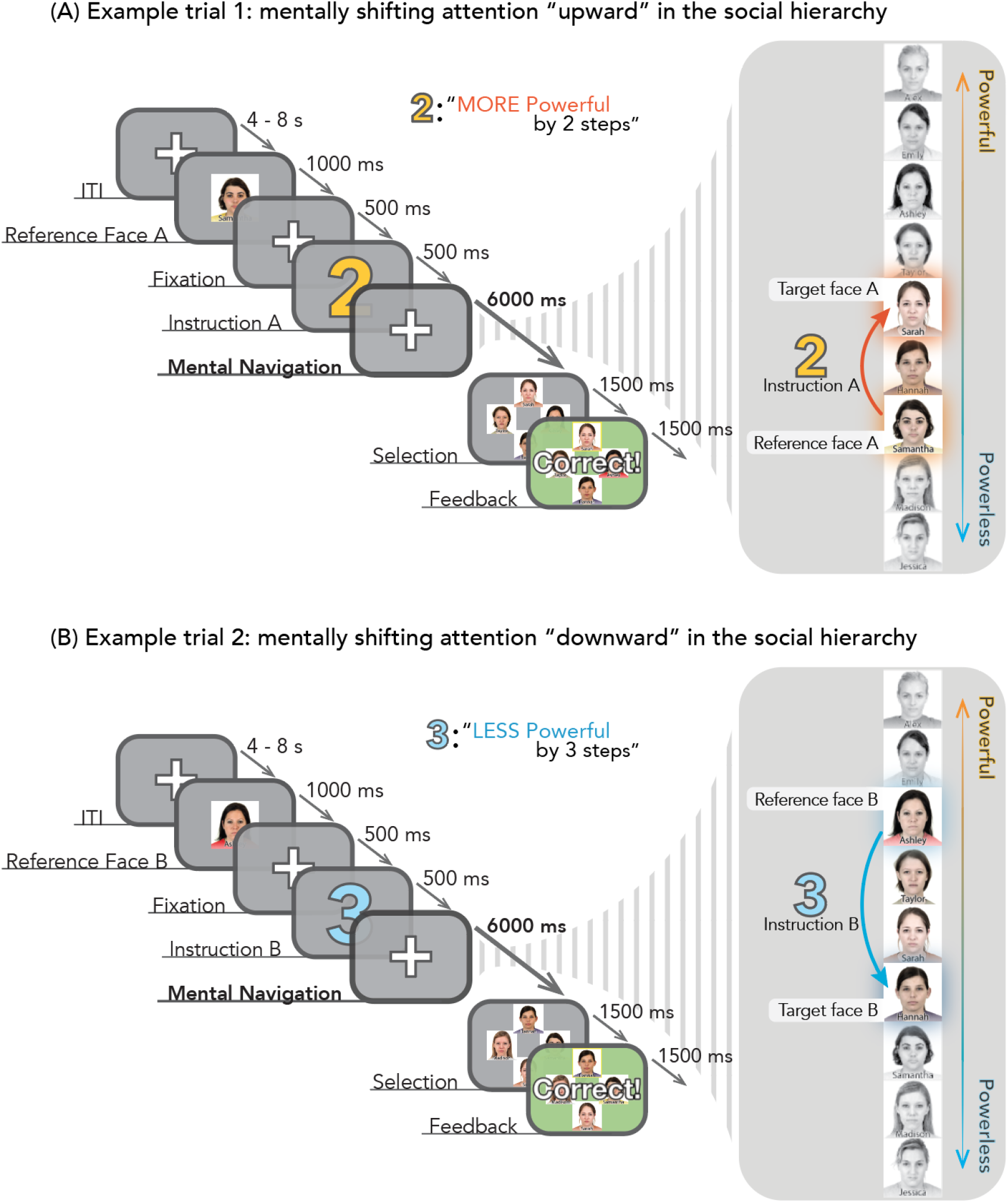
Mentally “navigating” the hierarchy (Task II). In each trial, participants were shown a reference face, followed by a colored number. The color of the number indicated whether they would need to determine the identity of a person more (**A**) or less (**B**) powerful than the reference person. The number itself indicated the number of steps participants would need to move from the reference person to reach the target person. Next, participants had 6 seconds to mentally “navigate” through the hierarchy and identify the target person. They were then prompted to select an answer from four options, and feedback was presented afterwards. The response options were displayed very briefly to force participants to arrive at their answers during the Mental Navigation portion of the trial.

**Figure 4.**
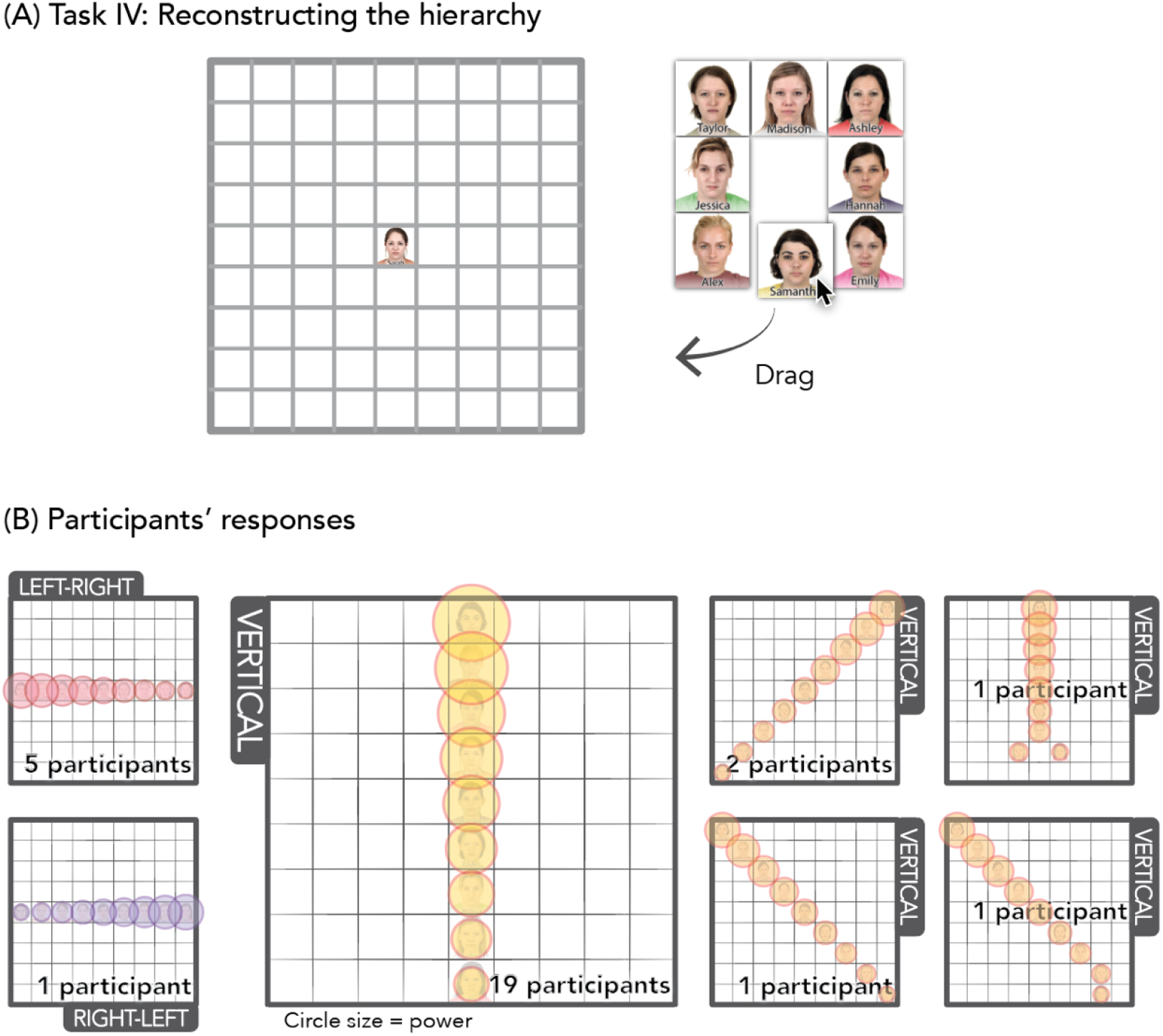
Methods and results for hierarchy reconstruction task (Task IV). (**A**) In this task, participants arranged the faces on a 9×9 grid, in any configuration that they thought best represented these people’s relative power. The middle person (i.e., the 5th person) in the hierarchy was already set at the center, and participants were asked to drag and drop the other 8 faces onto the grid. (**B**) All participants made linear arrangements, or close to linear arrangements, which all accurately reflected the relative status of faces. Based on the overall structure, 20 of the 30 participants arranged the social hierarchy in a top (most powerful) to bottom (least powerful) manner, and 10 participants arranged them left-to-right (*n* = 5), diagonally (*n* = 4), or right-to-left (*n* = 1). In cross-task analyses of fMRI data, we related multivoxel response patterns during social hierarchy navigation to vertical shifts of spatial attention if a participant responded with a vertical or diagonal representation^1^ here (i.e., more power corresponds to higher locations in space), and to horizontal shifts of spatial attention if they arranged the faces horizontally here (i.e., more power corresponds to more leftward or rightward locations in space, depending on whether a participant arranged faces in a left-right or right-left configuration, respectively).

## Results

### Behavioral Data

#### Learning the Social Hierarchy

Participants first learned about the structure of a fictive social hierarchy of 9 people through trial and error, using a task adapted from prior related work (Kumaran et al., 2012) (Fig. 2). Participants were told that we were interested in how people learn about social information and that they would be learning about who had more power in an organization. On each trial, participants chose which of two people they thought had more power in the organization. All phases of this task were designed to avoid biasing participants towards any particular mapping between locations in space and in the social hierarchy, as well as more generally, between the social and spatial domains. For example, faces were presented one at a time in the same location on the screen (i.e., faces were never shown above or beside one another). Participants in the fMRI study attained an average accuracy of 98.24% (*SD* = 1.67%) in the last 3 blocks of Session 1, indicating that they had effectively learned the social hierarchy.

#### Shifting Attention in Social Hierarchy Knowledge and in External Space

After learning the social hierarchy, participants performed a second task involving mentally “navigating” (shifting attention within) knowledge of the social hierarchy (Fig. 3; Methods). On each trial, participants saw a reference face, then were instructed to mentally “navigate” towards a more or less powerful person in the hierarchy. Participants who successfully performed this task in Session 1 (see Methods for more details) were eligible to return for the fMRI study (Session 2; see Fig. 1). During the fMRI study, participants completed several rounds of the social hierarchy navigation task (average accuracy = 94.35%, *SD* = 3.08%). Participants then performed a second fMRI task (adapted from previous work; Knops et al., 2009) involving overt shifts of visual attention upward, downward, leftward, and rightward in space (Fig. 1; Methods). Together, these two tasks allowed us to characterize the neural response patterns associated with shifts of attention in different directions in internally represented knowledge of the social hierarchy and in external space.

#### Probing Mental Representations of the Social Hierarchy

After scanning, participants completed tasks probing their mental representation of the social hierarchy. Participants performed a drag-and-drop task where they arranged hierarchy members’ faces into whatever configuration they thought best represented their relative power in the organization (Fig. 4a; Methods). All participants’ relative ordering of faces matched the actual status order of faces. The majority of participants (*n* = 20) arranged faces vertically, with the most powerful people at the top, which is consistent with everyday language usage such as “top of the pecking order” and “low status”; about one third of participants used alternative (i.e., diagonal, *n* = 4; horizontal, *n* = 6) configurations (Fig. 4b). Each participant’s response on this spatial arrangement task informed how correspondences between the neural encoding of shifts of attention in space and in social knowledge were tested in cross-domain fMRI analyses (see Fig. 4b).

### Neuroimaging Data

#### SPL Encodes the Direction of Attentional Shifts in Social Knowledge

We first tested if the SPL encoded the “direction” of shifts of attention within knowledge of social relations. To do this, we used data from the social hierarchy navigation task (see Fig. 3) to test if multivoxel response patterns evoked by attentional shifts within knowledge of the social hierarchy in the same “direction” (e.g., two trials where the participant had to determine who was “above” a reference person in the social hierarchy) would be more similar than those evoked by attentional shifts in opposite “directions” in knowledge of the social hierarchy. To quantify this, we computed an average pattern similarity score (*Sim*) between multivoxel response patterns evoked by shifts of attention in the same directions (*Sim_matching_*) and in opposite directions (*Sim_mismatching_*), then compared these two values (see Fig. 5a and Methods). As shown in Fig. 5a, multivoxel response patterns in both the left and right SPL signaled directions of attentional shifts in knowledge of social relations. In other words, *Sim_matching_* was significantly greater than *Sim_mismatching_* in both the left (*t*(29) = 5.373, *p* = 4.497 × 10^−6^, *d_z_* = 0.981) and right (*t*(29) = 4.300, *p* = 8.804 × 10^−5^, *d_z_* = 0.785) SPL (unless otherwise indicated, all reported *p*-values are one-tailed, given that all hypotheses were directional; effect sizes were calculated using Cohen’s measure of effect size for paired samples, *d_z_* ^34^). That is, when mentally navigating the social hierarchy, the SPL encoded attentional shifts in the same direction with more similar multivoxel patterns, compared to attentional shifts in different directions.

**Figure 5.**
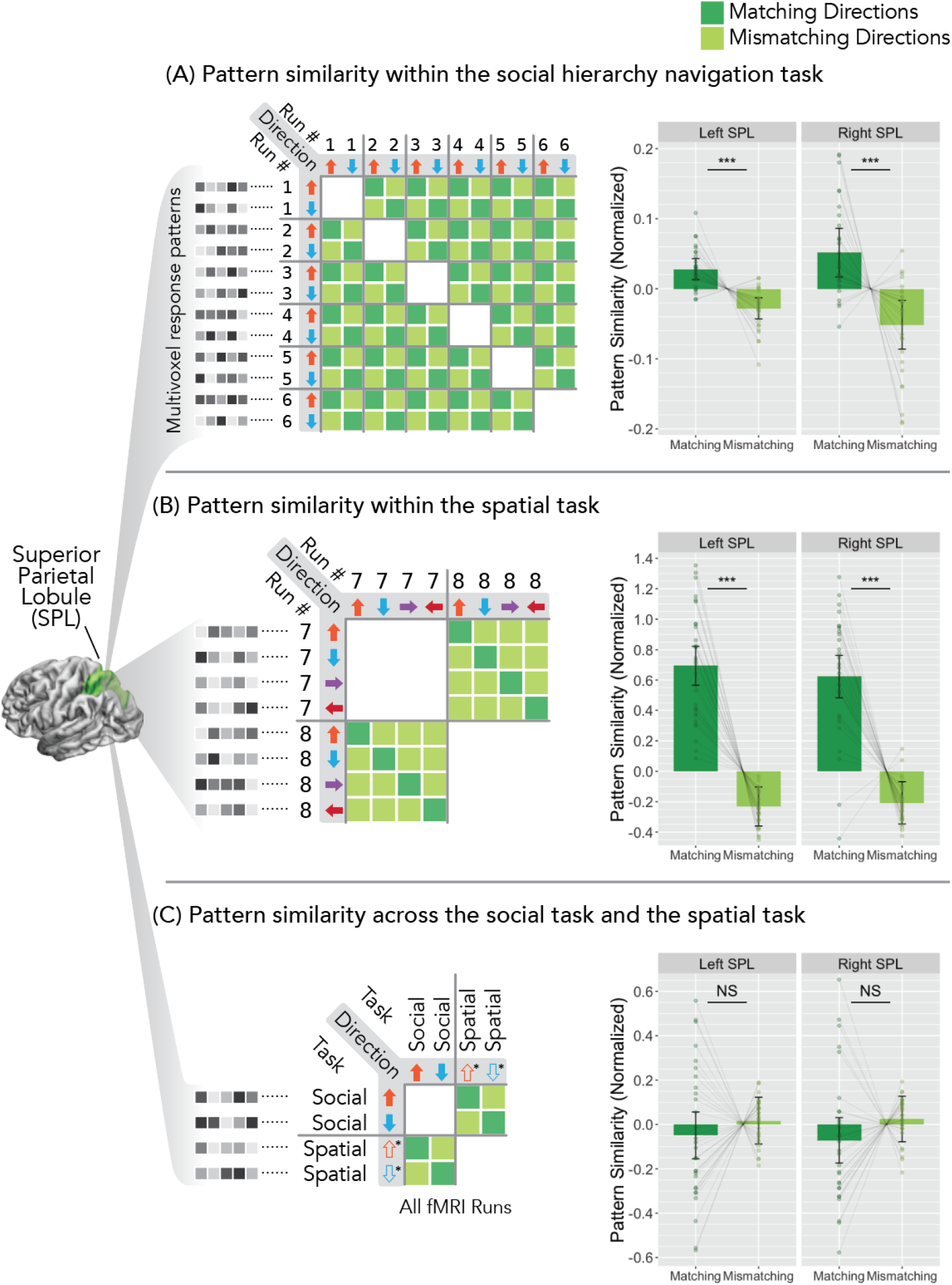
The SPL signals directions of attentional shifts in external space and within internal representations of social relations in overlapping but distinct multivoxel response patterns. **(A)** Average multivoxel response patterns in the SPL were calculated for shifts of attention “upward” and “downward” within knowledge of the social hierarchy for each participant. Mean similarities of multivoxel response patterns in the SPL across runs (*Sim*) were computed for each participant for matching (dark green) and mismatching (light green) directions of attentional shifts in social hierarchy knowledge. In the social task, multivoxel response patterns corresponding to matching directions of attentional shifts were significantly more similar to each other than those for mismatching directions, in both the left and right SPL. The same data analytic procedure was repeated to compare cross-run pattern similarities (*Sim*) in the SPL for matching (dark green) and mismatching (light green) directions of attentional shifts in the spatial attention task (**B**) and across social and spatial tasks (**C**). In the spatial attention task (**B**), multivoxel response patterns for matching directions of attentional shifts were more similar to each other than those for mismatching directions of attentional shifts in both the left and right SPL. When comparing multivoxel response patterns for corresponding “directions” across the social and spatial tasks (**C**), response patterns for matching directions (based on each participant’s responses to the social hierarchy reconstruction task) were not more similar than those for mismatching directions in either the left or right SPL. Further, results of cross-task Bayesian hypothesis tests were also in favor of the null hypothesis that pattern similarity scores were the same when comparing attentional shifts in matching and mismatching directions in space and social knowledge (*Sim_matching_* = *Sim_mismatching_*). Thus, multivoxel response patterns in the SPL encoded the direction of attentional shifts in external space (**B**) and within internal representations of social relations (**A**), but attentional shifts in matching directions were not signaled by the same patterns across social and spatial tasks (**C**). Error bars show 95% CIs. \*\*\**p* < .0001

#### SPL Encodes the Direction of Attentional Shifts in External Space

We next used analogous data analytic methods to test for the encoding of the directions of overt attentional shifts in space (Fig. 5b). The SPL also encoded shifts of attention in external space: Spatial attentional shifts in matching directions evoked more similar multivoxel response patterns than those in mismatching directions, in both the left (*t*(29) = 10.413, *p* = 1.306 × 10^−11^, *d_z_* = 1.901) and right (*t*(29) = 8.648, *p* = 7.995 × 10^−10^, *d_z_* = 1.579) SPL.

#### Does the SPL Encode Directions of Attentional Shifts in Social Knowledge and External Space in the Same Way?

Given that the directions of attentional shifts in external space and internal representations of social relations were both encoded in multivoxel response patterns in the SPL, we next tested if the SPL encoded shifts of attention in space and social knowledge in the same way. To determine which direction of spatial attentional shifts (up, down, left, right) corresponded to which direction of attentional shifts in social knowledge (more powerful, less powerful) for each participant, we used the mappings between locations in space and the social hierarchy that the participant indicated in the post-scan drag-and-drop task (Fig. 4). For example, if, after scanning, a participant had arranged the faces vertically, with the most powerful people at the top, we tested if neural response patterns evoked when shifting attention towards more powerful people in the social hierarchy resembled those evoked when shifting attention upward in space. On the other hand, if a participant had arranged the faces horizontally, with the most powerful people on the left, we tested if neural response patterns evoked when shifting attention towards more powerful people resembled those evoked when shifting attention leftward in space. As shown in Fig. 5c, in this cross-domain analysis, attentional shifts in matching directions across domains were not more similar than attentional shifts in mismatching directions in either the left (*t*(29) = −0.911, *p* = 0.815) or right (*t*(29) = −1.354, *p* = 0.907) SPL. Furthermore, we conducted a Bayesian hypothesis test to compare the marginal likelihoods of the null model *Sim_matching_* = *Sim_mismatching_* (i.e., multivoxel response pattern similarities were equivalent for attentional shifts in matching and mismatching directions across the social and spatial domains), versus the alternative model *Sim_matching_* ≠ *Sim_mismatching_* (i.e., multivoxel response pattern similarities were different when attentional shift directions were matched and when they were mismatched across the social and spatial domains). The resulting Bayes factors were moderately in favor of the null model in the left SPL (BF_10_ = 0.284), and weakly in favor of the null model in the right SPL (BF_10_ = 0.444). Taken together, these results suggest that overlapping but distinct codes in the SPL encode shifts of attention in social knowledge and in external space.

### Do Other Brain Regions Encode Shifts of Attention in Social Knowledge and External Space? Are They Encoded in the Same Way?

We next conducted exploratory analyses testing if other brain regions encode shifts of attention in external space and knowledge of social relations, and if they are encoded in the same way. These analyses were conducted in two ways. First, we repeated the pattern similarity analyses that had been conducted in the SPL in anatomically defined parcels spanning all of cerebral cortex and subcortical gray matter structures (see Methods; Figs. 6a and 7). Second, we performed a whole-brain searchlight analysis (see Methods; Fig. 6b). As shown in Fig. 6, results from the parcel-based and searchlight approaches were largely consistent with one another.

**Figure 6.**
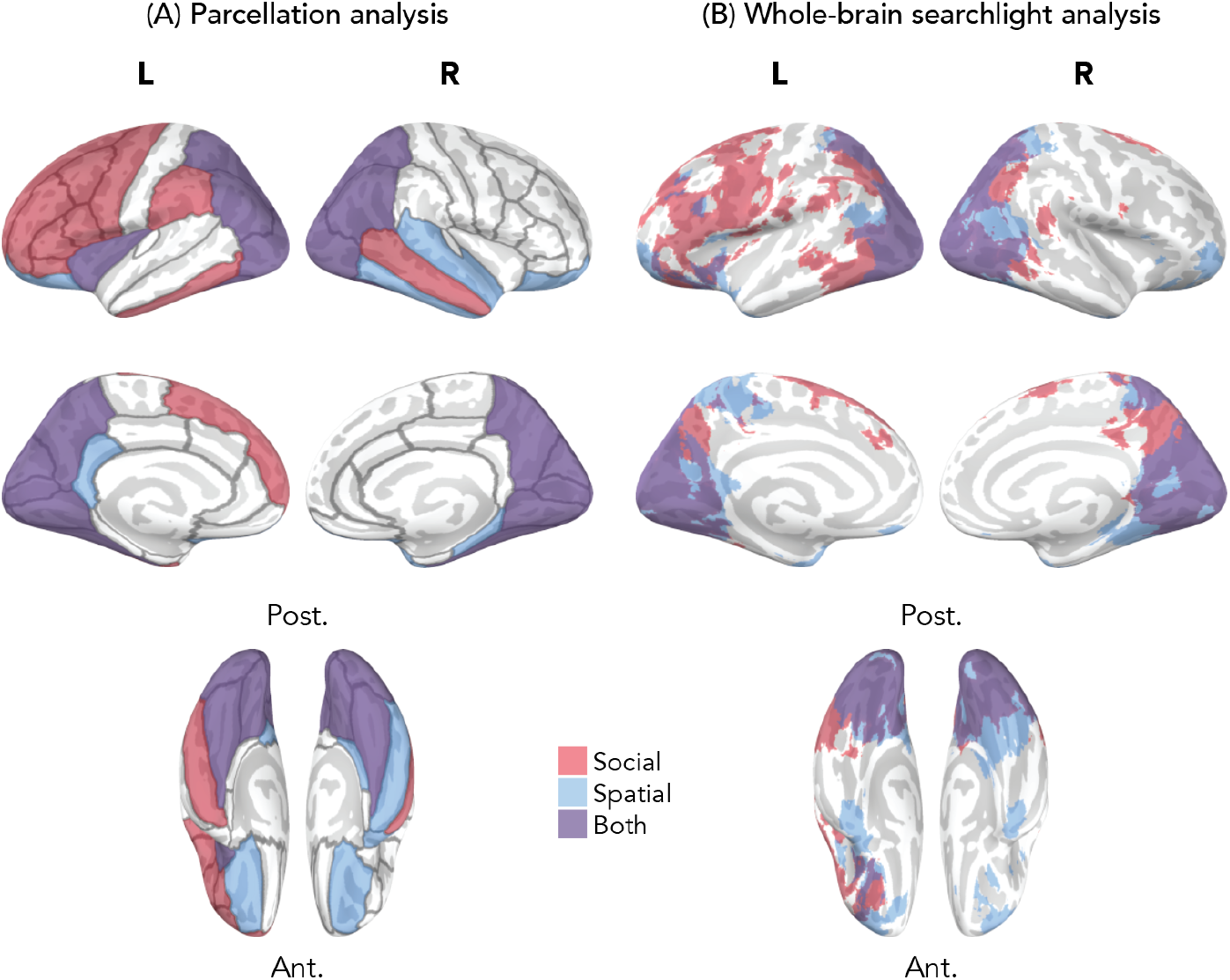
Shifting attention in space and social knowledge appears to be supported by distinct codes carried in largely overlapping brain regions. Both panels show results of exploratory whole-brain pattern similarity analyses. Red indicates areas that encoded “directions” of attentional shifts in social knowledge, blue indicates areas that encoded directions of attentional shifts in the spatial attention task, and purple indicates areas that encoded “directions” of attentional shifts in both tasks. A similar pattern of results was observed when analyses were conducted within anatomically defined parcels **(A)** and with a searchlight approach **(B)**. Cross-domain pattern similarity analysis did not reveal any brain regions that encoded attentional shifts similarly across tasks. All visualized results are significant at a corrected threshold of *p* < .01 (one-tailed).

**Figure 7.**
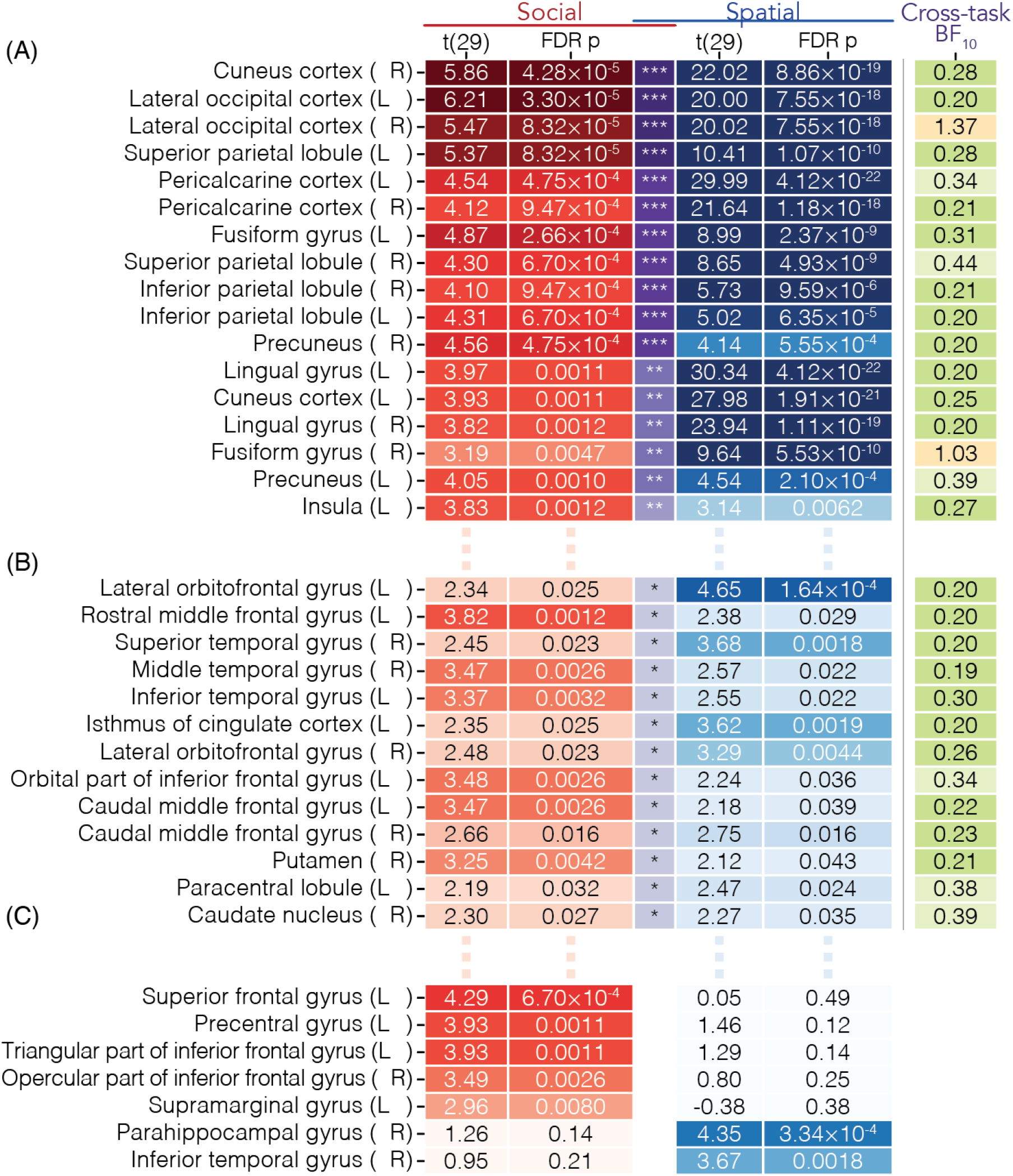
Regions that encode the direction of attentional shifts in space, social knowledge, or both: Results of exploratory whole-brain pattern similarity analyses. This table shows brain regions where within each task, multivoxel response patterns for matching directions of attentional shifts were significantly more similar to each other compared to those for mismatching directions (i.e., this figure provides additional details for the results shown in Fig. 6a). The upper two panels, (**A**) and (**B**), contain brain regions where multivoxel response patterns encoded the direction of attentional shifts (i.e., where *Sim_matching_* > *Sim_mismatching_*) in both social and spatial tasks at significance levels of *p* < .01 and *p* < .05, respectively (FDR-corrected). Asterisks and shades of purple in the middle column show the significance levels that *p*-values met in both social and spatial tasks: \**p* < .05 (FDR-corrected) on both tasks, \*\**p* < .01 (FDR-corrected) on both tasks, \*\*\**p* < .001 (FDR-corrected) on both tasks. Areas in occipital and parietal cortex most robustly encoded attentional shifts in both social and spatial tasks. That said, cross-domain pattern similarity analyses suggested that no regions encoded shifts of attention in corresponding “directions” in social knowledge and external space in the same way. Additional Bayesian hypothesis tests found weak (0.33 < BF_10_ < 1) to moderate (0.1 < BF_10_ < 0.33) evidence that most of these brain areas did not encode attentional shifts in social and spatial tasks in the same way (green cells); except that in the right lateral occipital cortex and right fusiform gyrus (yellow cells), there was weak evidence (1 < BF_10_ < 3) favoring the opposite. The lower panel (**C**) contains brain regions that encode attentional shift directions in either the social or spatial task (thresholded at *p* < .01), but not both. Comprehensive results for all brain regions are shown in Figure 7–figure supplement 1.

#### Exploratory Whole-Brain Pattern Similarity Analyses

Results showed considerable overlap in the brain regions that encoded the direction of shifts of attention in both tasks, particularly in regions of parietal and occipital cortex (Figs. 6 and 7ab). Additionally, several brain regions encoded the direction of attentional shifts only in social knowledge (e.g., left lateral prefrontal cortex, bilateral supramarginal gyri; Fig. 7c) or only in external space (e.g., hippocampus, parahippocampal cortex, entorhinal cortex; Fig. 7c).

Given the considerably overlapping encodings across social and spatial tasks in the parietal and occipital cortex, we further examined whether these (or any) regions encoded the direction of attentional shifts in the two domains in the same way. However, exploratory whole-brain analyses did not identify any brain regions where matching directions of attentional shifts in social knowledge and external space were encoded similarly. Additional Bayesian hypothesis tests were conducted to assess the performance of a null model (*Sim_matching_* = *Sim_mismatching_*) versus an alternative model (*Sim_matching_* ≠ *Sim_mismatching_*) in the brain regions where both social and spatial shifts of attention were encoded; the results are shown in Fig. 7 (“cross-task BF_10_”). Similar to our primary SPL-focused analyses, these results suggested that while there is much overlap in the brain regions that encode shifts of attention in space and social knowledge, these mental operations may be signaled by distinct neural population codes in most implicated brain regions.

#### Exploratory Whole-Brain Cross-Domain Classification Analyses

To further test the above conclusion from pattern similarity analyses, we performed cross-domain classification analyses, both within the SPL and across the brain. As pattern similarity analyses are based on correlations of all voxels in a region, they weight values from each voxel in a pattern equally. In contrast, methods such as cross-domain decoding using support vector machine (SVM) learning are able to assess if encoding mechanisms are shared across domains (Chavez et al., 2017; Parkinson et al., 2014) in a manner that gives higher weights to more relevant voxels (Haynes, 2015). Thus, we performed cross-domain decoding analyses using SVM classification with both the parcel-based and searchlight approaches. SVM classifiers were trained to decode directions of spatial attentional shifts based on evoked neural response patterns, then tested on their ability to decode the direction of attentional shifts in social knowledge, and vice versa. Again, participants’ responses on the post-scan drag-and-drop task determined which directions in space were matched to which “directions” in the social hierarchy. Results showed that in parcel-based and searchlight analyses, there were no regions where cross-domain decoding was possible. Additional exploratory analyses employing hyperparameter tuning and other classification algorithms (e.g., logistic regression), also yielded null results (see Supplementary file 2). Follow-up analyses indicated that for each task, within-task decoding of the direction of attentional shifts was possible in many of the same regions of parietal and occipital cortex that were implicated in the within-task pattern similarity analyses (see Supplementary file 1). This suggests that the lack of success in cross-domain decoding in these regions did not result from an inability of the algorithms to learn distinctions within each task, but may instead stem from the distinctions learned within tasks not generalizing across tasks. Taken together, these results cohere with those of our primary analyses (Fig. 5), and suggest that regions of parietal and occipital cortex encode the direction of attentional shifts in both social knowledge and external space, but this information is likely encoded using distinct underlying mechanisms.

## Discussion

Here, we found that the SPL, a region with a long-established role in directing shifts of attention in external space, also encodes shifts of attention in internal representations of social relations. Multivoxel response patterns in this region signaled whether participants were mentally shifting attention “upward” or “downward” in knowledge of a social hierarchy (i.e., towards more powerful or less powerful people). Convergent results were obtained using both pattern similarity and decoding analyses. In line with prior work, multivoxel response patterns in the SPL also encoded the direction of shifts of attention in external space. Furthermore, exploratory whole-brain analyses implicated additional regions of the posterior parietal cortex and occipital cortex in encoding direction of attentional shifts in space and social knowledge.

Thus, shifts of attention in internal representations of social knowledge and in external space are encoded by partially shared brain regions. At the same time, in these implicated brain regions (including SPL), cross-domain pattern similarity and decoding analyses illustrated that shifts of attention in space and social knowledge may be encoded in functionally independent response patterns. These results shed light on the neural basis of reasoning about social relations, and suggest that shifts of attention in space and in social knowledge are supported by distinct neural mechanisms within largely overlapping brain regions.

### Multivoxel Patterns in the SPL Encode Shifts of Attention in Social Knowledge

Pattern similarity analyses were used to characterize the mental operations within the SPL that are involved in reasoning about social relations. This data analytic approach was chosen so that we could rule out the possibility that SPL was merely recruited when processing social hierarchy knowledge without actually encoding mental operations within such knowledge. For example, SPL responses could have been organized only by task difficulty (e.g., with pattern similarity and/or response magnitude only reflecting the similarity of the amount of “distance traveled” on particular trials^2^). That possibility would have been consistent with prior work where more difficult social status comparisons evoked stronger responses in the vicinity of this region (Chiao et al., 2009). However, we found that the multivoxel pattern similarity in the SPL reflected the “direction” in which people were shifting attention in their knowledge of the social hierarchy, suggesting that the SPL directly encodes the attentional shifts in social knowledge.

### Why Would the SPL Encode Shifts of Attention in Space and Social Knowledge?

The SPL has been widely implicated in shifting attention in the external world (Molenberghs et al., 2007; Serences et al., 2004). The fact that it also encoded the “direction” of attentional shifts within social knowledge is consistent with behavioral and linguistic evidence that operating on space and social knowledge may involve common representational resources (Dai and Zhu, 2018; Schubert, 2005; Zanolie et al., 2012). Furthermore, it has been suggested that over the course of human brain evolution, regions of the PPC expanded (Van Essen et al., 2001) and formed new connections (Mantini et al., 2013) as they came to support increasingly abstract aspects of social cognition (Parkinson and Wheatley, 2013; Yamazaki et al., 2009). Thus, as recently suggested by Bottini and Doeller (2020), the SPL may support attentional shifts in internal map-like representations of relational knowledge, in spatial, social, and other domains (this knowledge may be stored in hippocampal-entorhinal cortex – Garvert et al., 2017; Tavares et al., 2015). This is consistent with the substantial overlap we found in the brain regions that encoded spatial and social shifts of attention (Figs. 6 and 7a), which could reflect similarities in the processes involved in these mental operations. In other words, shifts of attention in social knowledge may be supported by brain regions with a long-standing role in spatial attention because characteristics of such brain regions (e.g., their internal structure and connectivity with other brain regions) afford operations common to directing shifts of attention in space and in knowledge of social relations (e.g., coordinate transformations).

### Shifting Attention in Space and Social Knowledge is Supported by Distinct Codes within Overlapping Brain Regions

Relatively little is known about how the brain supports mental navigation of social relations. Further, most relevant past research has used mass univariate analyses, and thus, would be insensitive to effects carried in multivoxel response patterns. Therefore, we performed exploratory whole-brain analyses in addition to our main analyses on the SPL. The cross-domain pattern similarity analyses found several regions that encode shifts of attention in both space and social knowledge, including superior and inferior aspects of lateral PPC, as well as the precuneus, fusiform cortex, and regions of occipital cortex (Figs. 6 and 7a).

Considerable efforts were taken to avoid biasing participants toward thinking about the hierarchy spatially. For example, instructions avoided the use of spatial language and faces were never shown above, below or beside one another during the learning task (Fig. 2). In the response selection phase of the social task, faces (Fig. 3) and response button choices were configured such that there was no systematic relationship between response location–on the screen or response pad–and faces’ positions in the hierarchy. Additionally, participants were not informed of the spatial attention task until after all social task trials had been completed to avoid inducing spurious similarities between how participants would perform the “social hierarchy navigation” task and the spatial attention task. Further, participants were instructed to focus on the center of the screen during the social task, and the results of an additional eye-tracking study (see Supplementary file 3) suggest that neural activity during the social task did not reflect eye movements. Still, there was considerable overlap in the brain regions encoding social and spatial operations. As discussed earlier, this is consistent with previous behavioral research linking mental representations of space and social hierarchy knowledge (e.g., Dai and Zhu, 2018; Schubert, 2005; Zanolie et al., 2012). Moreover, participants’ post-scan responses indicated that they tended to ascribe a systematic spatial organization to people’s relative locations in the status hierarchy (usually a vertical organization, see Fig. 4), suggesting an association between spatial and social contents in their internal representations.

At the same time, brain regions encoding both social and spatial mental operations appeared to encode them in largely independent coding schemes. Using both anatomically defined parcels and searchlights, the cross-domain pattern similarity analyses did not identify any region that encoded attentional shifts in social knowledge and space in the same way. As described in the Results section, we complemented these cross-domain pattern similarity analyses with exploratory decoding analyses to provide sensitivity to distinctions driven by only a subset of voxels within a region. Whole-brain cross-domain decoding analyses conducted within both anatomically defined parcels and searchlights identified no region where directions of attentional shifts in space and social knowledge were encoded similarly, consistent with findings from whole-brain pattern similarity analyses. Thus, despite considerable overlap in the brain regions involved in shifting attention in space and social status knowledge, we did not find evidence for a common encoding of these two mental operations. Notably, this was not because these analyses lacked sensitivity to within-domain information: Both pattern similarity analyses and classification analyses consistently identified regions, including regions with long-established roles in spatial attention, that signaled directions of attentional shifts within each domain (see Fig. 6 and Supplementary file 1). Nonetheless, these reliable within-domain distinctions did not tend to generalize across domains. Taken together with the results of Bayesian hypothesis tests, these results suggest that shifting attention in space and social knowledge relies on overlapping but largely distinct coding schemes. As mentioned above, encoding shifts of attention in abstract knowledge of social relations may take place in regions that also encode shifts of attention in space because of partially shared processing demands for these mental operations.

Several psychological factors may have contributed to neural response patterns in the social task being distinct from those in the spatial task. For example, it is possible that participants’ post-scan arrangements of the social hierarchy, which we used to inform the cross-domain correspondence of directions in our analyses, do not accurately reflect participants’ actual mental representations during the social task. Rather, their actual representations may have been hard to draw or may have changed over time, or it may be difficult for participants to accurately introspect about how these mental representations are organized. Alternatively, although the SPL encodes directions of attentional shifts in both social and spatial domains, it may encode the operation of shifting attention upward in space completely differently from shifting attention “upward” abstractly in social hierarchy knowledge. In other words, the two corresponding “directions” of attentional shifts across domains may be represented distinctly by the brain, even though most participants mapped “higher” positions in the hierarchy to higher positions in space (Fig. 4), and even though we describe both as “upward” in common parlance. That is to say, “directions” within representational spaces encoded by the SPL (and other brain regions) may not have straightforward correspondence across the two domains. Additionally, it is also possible that aspects of the tasks used may have contributed to evidence for distinct coding schemes for shifts of attention in space and social knowledge, as discussed in the subsequent section.

### Limitations and Future Directions

The current study focused on a single kind of social relational knowledge (a status hierarchy). This type of knowledge afforded a clear hypothesis for the nature of cross-domain mappings, which was necessary to examine if any brain regions encode shifts of attention in space and social knowledge in the same way. Thus, in this study, social hierarchy knowledge served as a practical place to start for examining these questions. Future research can extend these findings by examining how the brain supports mental navigation of other kinds of social relational knowledge, and how this compares to other kinds of mental operations.

We also used very constrained social tasks. Using a highly controlled social task allowed us to ascertain when participants were mentally traveling “up” or “down” in their knowledge of the social hierarchy, which was necessary to extract corresponding neural response patterns. Furthermore, a few considerations suggest the efficacy of the hierarchy learning task in eliciting social processing. First, this task was adapted from a prior study (Kumaran et al., 2012), in which providing a social context merely through verbal instructions evoked distinct neural processes for learning social, compared to non-social, hierarchies. Second, this task resembles real-world social status hierarchy learning in that in both cases, status knowledge largely emerges from observing relations between pairs of individuals and using transitive inference to construct mental representations of social hierarchies (Cheney and Seyfarth, 1990a, 1990b). However, both social tasks differed in many respects from how social relations are learned and navigated in the real-world. Other researchers (Tavares et al., 2015) have developed ecologically valid paradigms to examine how the brain tracks subjective affiliation and power during social interactions. Future research could extend the current work using similarly ecologically valid paradigms, such as naturalistic social interactions or veridical social knowledge.

Future research may also further address whether shifts of attention in social knowledge and space are encoded in the same way by employing different study designs. First, we chose social and spatial tasks that were very different from each other in order to establish the validity of our primary results that showed overlapping encodings for attentional shifts in space and social knowledge. If the two tasks were similar to each other, the overlapping results would have been confounded by the similarities between tasks themselves. However, the distinctness of the two tasks may also have been a factor that made cross-domain decoding difficult. Future studies may match tasks from different domains more closely, for example, in terms of task difficulty, dimensionality, and self-versus world-centeredness, which may influence neural encodings of cognitive maps (Bottini and Doeller, 2020). Second, although the results of analyses using Bayes factors were somewhat in favor of the null hypothesis in most brain regions (i.e., most brain regions may not encode attentional shifts in social knowledge and space in the same way), the evidence was not strong. Future research may further investigate this question using additional analytical methods and paradigms.

It would also be interesting to compare the neural response patterns evoked when mentally navigating social and non-social relational knowledge (e.g., shifting attention between memories of different spatial locations, rather than between different spatial locations themselves; other forms of relational knowledge). Past research has shown that shifting attention along the mental number line (as in mental arithmetic) evokes patterns similar to shifting attention in external space (Knops et al., 2009). The asymmetry between those results and the results of the current study suggests that shifts of attention in knowledge of social relations and other forms of relations (e.g., numerical magnitude) would be encoded differently, but future studies could more directly tease these phenomena apart. Regardless of the results of such studies, the capacity to perform transitive inferences on relational knowledge may be fundamentally social in origin: This capacity is thought to be particularly important for highly social species living in large groups containing linear status hierarchies (Cheney and Seyfarth, 1990a; Hogue et al., 1996) and may have evolved to support social complexity (Maclean et al., 2008).

## Conclusions

Humans have a sophisticated capacity to learn and reason about social relations. The current results demonstrate considerable overlap in the brain regions that encode shifts of attention in the space around oneself and in internal representations of social knowledge. At the same time, within these overlapping brain regions, multivoxel codes within each domain (space, social knowledge) were largely independent. These findings provide insight into the neural basis of reasoning about knowledge of social relations. More generally, these findings also highlight the value of examining distributed response patterns for elucidating the diverse mental operations that brain regions encode, and how they relate to one another.

## Supporting information

Supplementary file 1. Within-domain decoding

Supplementary file 2. Hyperparameter tuning for cross-domain classification analysis

Supplementary file 3. Eye-tracking study

Supplementary file 4. Reference face analysis

Supplementary file 5. Alternative analyses for participants with diagonal representations of the social hierarchy

Supplementary file 6. Analysis of the magnitude of attentional shifts within social knowledge

## Acknowledgments

The authors wish to thank Ryan Hyon, Savanna Gharibian, and Georgia Perris for assistance with data collection, as well as those who provided helpful feedback on the initial presentation of these results at the 2018 meeting of the Social and Affective Neuroscience Society. This work was supported by funding from a UCLA Academic Senate Council on Research Faculty Research Grant, a Sloan Foundation Research Fellowship, and the UCLA Center for Cognitive Neuroscience.

## Author Contributions

Conceptualization, C.P.; Methodology, C.P. and M.D.; Software, M.D.; Formal Analysis, M.D.; Investigation, M.D. and R.B.; Resources, C.P.; Data curation, M.D.; Writing - Original Draft, C.P. and M.D.; Writing - Review & Editing, C.P., M.D., and R.B., Visualization, M.D.; Supervision, C.P.; Project Administration, C.P.; Funding Acquisition, C.P.

## Data and Code Availability

Code and de-identified behavioral data have been made publicly available and are accessible at https://github.com/CSNLab/hierarchy-study.

## Materials and Methods

### Overview of Paradigm and Analysis

The overall procedure of this study is illustrated in Fig. 1, and is outlined below and in the Results section. Detailed descriptions of each component of the study are provided in the Procedure section. Participants attended a behavioral session (Session 1) and a subsequent fMRI session (Session 2). Qualifying participants completed the fMRI session 2.5 hours − 5.85 days after the initial behavioral session (*M* = 2.71 days).

In Session 1, participants first learned a linear social hierarchy of 9 people through trial and error (Task I, adapted from a prior study; Kumaran et al., 2012), and then performed a social hierarchy navigation task (Task II), where they were repeatedly asked to shift attention within their knowledge of the social hierarchy (i.e., to find a target person in the hierarchy, given another reference person, a distance and “direction”). Participants who had satisfactory accuracy (>70%) in Task II of Session 1 were invited to participate in Session 2, which involved performing the social task (Task II) again in the MRI scanner.

Upon arriving for Session 2, participants first completed short versions of Tasks I and II again outside of the scanner to refresh their memories of the social hierarchy and of the social task, respectively. Then, while being scanned, participants performed the same social task (Task II), followed by a task involving overt shifts of attention (i.e., eye movements) in different directions in external space (Task III, adapted from a prior study; Knops et al., 2009). After scanning, participants’ mental representations of the social hierarchy were probed by asking them to rearrange the faces in whatever spatial configuration they thought best represented the people’s relative power (Task IV; Fig. 4). These responses were used to determine which directions of a participant’s attentional shifts should be matched across domains for each participant in pattern similarity analyses and cross-domain decoding.

Importantly, in the experimental materials, we avoided instructions, stimulus arrangements, and response formats that might lead participants to think about the social hierarchy in any particular spatial terms, or to visualize the hierarchy in any particular way. In the instructions, for example, we described the hierarchy as an “organization” where one person is a certain number of “degrees” more or less “powerful” than another person. During the entire experiment, faces from the hierarchy were displayed in one of three ways: a) alone at the center of the screen, or randomly arranged b) in a group of four shaped like a diamond (see the last two screens in Fig. 3), or c) in a group of 8 shaped like a square (a 3 × 3 grid with an empty central cell, see Fig. 4a). Thus, participants never saw the social hierarchy arranged in a linear configuration during the experiment; therefore, any linear arrangement of the hierarchy in their mental representations would have been derived on their own.

### Participants

Participants were recruited at the University of California, Los Angeles. 31 participants qualified for the fMRI study and completed all tasks after passing the accuracy criterion (70%) in Session 1, as described later in this section. Data from one participant were discarded from analysis due to excessive movements during the fMRI scan (>3 mm in all runs). All participants in the final sample were right-handed, native English speakers, and between the ages of 18 and 31 (*M_age_* = 20.17, 11 males, 19 females). Participants met standard MRI safety criteria, and provided informed consent in accordance with the policies of the UCLA Institutional Review Board.

### Stimuli

We obtained 18 face images (9 white males and 9 white females) from the Chicago Face Database (Ma et al., 2015) with approximately matched ratings of perceived age, affect, dominance, and attractiveness. The gender of the social hierarchy was matched to the gender of the participant. To facilitate participants’ ability to individuate and remember the people about whom they learned, and thus, reduce the difficulty of tasks, images were edited to make the T-shirts different colors (rather than all gray) and names were added below the faces. The names were selected from the most popular given names for babies born during the 1990s, as published by the US Social Security Administration (https://www.ssa.gov/oact/babynames/decades/names1990s.html). Assignment of faces to positions in the social hierarchy was randomized across participants.

### Procedure

All paradigms described below, except the social hierarchy reconstruction task (Task IV), were created using Python 2.7 with PsychoPy 1.85 (Peirce, 2007). The social hierarchy reconstruction task was created with HTML, CSS, and JavaScript.

#### Task I: Learning the social hierarchy

In the learning task (see Fig. 2, adapted from a prior study; Kumaran et al., 2012), participants learned each person’s relative position in the social hierarchy through trial and error. The following instructions were used to verbally inform the participants of the content of this task:

> *“Our experiment looks at how people learn social information. There will be two parts of the experiment in today’s session, and your performance in both of them will determine whether you are eligible for the second fMRI session. In the first part, you will be looking at pictures of nine people who all belong to an organization, and your goal will be to figure out who has more power in this organization and who has less power. You will see pictures of two people in a trial, and all you have to do is choose the person who has more power in the organization. You will be using trial and error… So you’ll have to guess a little bit in the beginning, and you will get a score that tells you if you chose right or wrong. It might seem a little difficult at first, but most people who keep trying do figure it out… So just try your best.”*

Participants were also informed that they would use the information they learned from this task in later parts of the experiment: *“When you’re done with this part of the experiment, you will use what you learn about the people from this part to complete the second part today and the rest of the experiment. That means your performance in the rest of the experiment depends on how well you learn the information from this part… So try to get the highest score possible in this part.”*

On each trial, participants saw two people and selected who had more power in the organization. Each trial started with a 1000 ms fixation cross, and then two faces were presented sequentially on the screen in a random order, with the presentation of each face lasting 1500 ms, with a 50 ms interval in between. Participants were then prompted to select the more powerful person in the pair by pressing and holding down the space bar, then releasing it when their preferred response choice appeared. Once the participant started to press the spacebar, the two face images were cycled through again in the same order in which they had been displayed at the start of the trial. While participants held down the spacebar, each face image was presented for 1000 ms at a time on the screen. An expanding circle was displayed behind each face; the size of this circle indicated how long the current face would remain on the screen. Participants then selected one of the faces by releasing the space bar when they saw the person they intended to choose. After a choice was made, the selected face was highlighted together with a score (e.g. “+10 points” after a correct answer or “-10 points” after an incorrect answer) displayed on the screen for 1000 ms. Afterwards, the two faces were presented again with the correct answers (i.e., who was relatively powerful and who was relatively powerless in the organization; see Fig. 2), and participants pressed “N” to advance to the next trial. During the entire task, stimuli and response options were not mapped to any spatial configuration (i.e., only one face was shown at a time at the center of the screen, instead of two faces appearing in a horizontal or vertical configuration; response choices between two options were made by releasing the spacebar when the preferred option appeared, rather than by pressing two response keys which would form a spatial configuration on the keyboard). This was to avoid inducing any extraneous mappings between positions in space and in the social hierarchy, and to avoid biasing participants towards thinking about the hierarchy in spatial terms any more than they typically would.

The task contained two types of trials, adjacent trials and non-adjacent trials. Each block in this task began with 16 adjacent trials in which the pairs of people presented were adjacent to each other in the social hierarchy (e.g., the most powerful person and the second most powerful person). Every possible adjacent pair of people was presented twice at the start of the block in a randomized order. The next portion of the block contained non-adjacent trials, in which the pairs of people presented were farther away from each other in the hierarchy (specifically, either two or three steps away from each other, e.g., the most powerful person and the third most powerful person). Thus, responding correctly on non-adjacent trials required participants to perform transitive inference on the social hierarchy knowledge that they had acquired during the first part of the block. During these non-adjacent trials, participants were also asked to rate how confident they were in their answer before receiving feedback about its accuracy. Each adjacent trial of a block was worth 10 points, each non-adjacent trial was worth 100 points, and participants were asked to try to get the highest score possible.

Participants first performed this task at the beginning of Session 1, in which a minimum of 12 blocks were displayed. If their accuracy was unsatisfactory after 12 blocks – i.e., they had either more than three errors in the adjacent trials of the last three blocks, or more than two errors in the non-adjacent portions of the last three blocks – the task would keep going until a maximum of 14 blocks had been completed.

At the beginning of Session 2, participants performed this task again outside of the scanner to refresh their memories of the social hierarchy. A minimum of four blocks were displayed during Session 2, and the task continued until either they had reached a perfect score in all of the last three blocks, or it had taken over 30 minutes.

#### Task II: Mentally navigating the social hierarchy

In this task, participants mentally shifted attention within the social hierarchy that they had learned in Task I in order to answer a series of questions (see Fig. 3). Each question asked the participant to find a target person in the hierarchy, given another person who served as a point of reference, a distance, and a direction. In each trial, the participant first saw the reference face from the hierarchy (1000 ms), and then a fixation cross (500 ms), followed by a colored number (500 ms). The number’s magnitude indicated the distance between the reference and the target person, and it was colored in either orange or blue, indicating the target person was either more or less powerful than the reference person. This mapping between color and direction was randomized across participants (i.e., for some participants, orange meant more powerful and blue meant less powerful; for the others, the opposite was true). Next, participants had 6000 ms to mentally “navigate” through their knowledge of the social hierarchy to determine their answers while a fixation cross was displayed. Afterwards, four faces were presented as response options, and the participants had 1500 ms to choose an answer. These faces were presented in a diamond configuration (Fig. 3) and while being scanned, participants responded using a response box with buttons configured in the same diamond shape (Current Designs 4-Button Diamond Fiber Optic Response Pad). Rather than using linear configurations of response choices and/or response buttons where faces and/or button options appeared side-by-side, we used this diamond configuration to avoid experimentally biasing participants to think more about the social hierarchy navigation task in any particular spatial manner. The correct answer was equally likely to appear in each of the four possible positions on the screen, and the incorrect response options were three other faces that were as close as possible to the correct answer in the hierarchy. After the response option screen had been presented, feedback was displayed for 1500 ms. If the participant did not make a choice in time, they were prompted to respond faster next time; otherwise, they were informed whether they were right or wrong while their selected answer was highlighted. Finally, a fixation cross was displayed between trials. During the behavioral session, each inter-trial interval (ITI) was 2 or 3 seconds. During the fMRI session, ITIs ranged from 4-8 seconds and were determined using NeuroDesign (Durnez et al., 2018), as described in more detail below.

Participants completed 3 blocks of this task at the end of Session 1, one block at the beginning of Session 2 right before they were scanned, and 6 runs inside the scanner. Each block contained 24 unique trials. For the blocks completed outside of the scanner, trial order was randomized. For those runs completed inside of the scanner, the stimuli order and ITI durations were generated using the Python package NeuroDesign (Durnez et al., 2018) to optimize our power to detect differences between attentional shifts in contrasting directions within each subject. The total duration was 402 seconds for 5 out of the 6 runs and 404 seconds for one run; run order was counterbalanced across participants using a Latin square design.

To ensure that participants were engaged in mental navigation during the 6000-ms period (which thus ensures the sensitivity of our fMRI analyses), we took efforts to minimize the possibility that they would be able to 1) predict the direction of the correct answer immediately upon seeing the reference face (and thus begin mental navigation away from the anchor face before the designated part of the trial), or 2) wait until after the response options appeared and then start figuring out the answer, or 3) reach an answer easily without “navigating” through the hierarchy. First, all of the reference faces and target faces were selected among the third to seventh people in the nine-person hierarchy, and the only possible distances between the two relevant faces were 2, 3, or 4 steps. Each of the 24 possible trials was presented once in each run. This avoided situations where a face from either extreme end of the hierarchy was used as the reference, in which case participants would have been able to ascertain the direction of navigation before seeing the instruction regarding direction (and thus begin mentally navigating away from the anchor face in that direction); it also avoided situations where participants could directly “jump” to an answer without needing to mentally navigate their knowledge of the hierarchy, e.g., when the answer was either around the top or bottom of the hierarchy, or adjacent to the reference face. Second, the four candidate response options were presented very briefly, so that participants were merely able to quickly skim the options and indicate the answer they had already figured out before seeing these options. This ensured that participants would have mentally navigated through their knowledge of the hierarchy before the options appeared and not after; if they did not already have an answer in mind before seeing the options, they would be unlikely to have enough time to figure it out when the options appeared. This aspect of the experimental design facilitated our ability to ascertain which time points reflected shifts of attention in particular directions in social hierarchy knowledge.

#### Task III: Shifting attention spatially

This task was adapted from previous related research (Knops et al., 2009). Participants were asked to follow a white target cross on the screen with their eyes. The target cross appeared at the center of the screen at the beginning of each trial, then moved up, down, left or right in 4 steps, and finally went back to the center during the ITI. Each step within a trial lasted for the same amount of time (500, 750 or 1000 ms), during which the target cross moved approximately 1.25° of visual angle, with a random jitter in the range of ± 0.42° in the direction of movement, and a random jitter of ± 0.26° perpendicular to the direction of movement. At the end of each trial, the target cross stayed in its last position for the first 3 seconds of the ITI to prevent inducing eye movements back to the center of the screen immediately after attentional shifts in the direction of interest. Then the target cross returned to the center and remained there for the second half of the ITI. The ITI durations were 6, 7 or 8 seconds. As in Task II, the order and timing of trials were determined using NeuroDesign (Durnez et al., 2018) in order to optimize power to detect differences between shifts of attention in different directions with each participant. Each run contained 40 trials and participants completed 2 runs, which lasted 409 seconds and 396 seconds.

#### Task IV: Reconstructing the social hierarchy

At the end of the experiment, we probed participants’ mental representations of the social hierarchy. Responses to the hierarchy reconstruction task (Task IV) were used to determine, for each individual participant, which directions of attentional shifts in the spatial task (Task III) corresponded to which directions of attentional shifts in the social task (Task II). In Task IV, we showed participants a 9 × 9 grid that was empty except for the central cell (see Fig. 4a). The middle person (i.e., the fifth person) in each participant’s social hierarchy appeared at the center of the grid, and the other 8 people appeared outside the grid in a random configuration. Participants were asked to drag the 8 people to the grid and arrange them in any configuration that they thought best represented these people’s relative power. Each participant’s answer was coded according to the way that they arranged the faces (see Fig. 4b).

### fMRI Data Acquisition

MRI data were collected using a Siemens 3 Tesla Prisma Fit MRI Scanner with a 32-channel head coil. Functional scans were obtained using a gradient echo sequence with 60 transverse interleaved slices (TR = 1000 ms, TE = 29 ms, flip angle = 52°, FOV = 198 mm, 2.4 mm isotropic voxels). For each subject, two field map scans were obtained before functional scans began in order to correct for the effects of field inhomogeneity. Participants used a 4-button diamond-shaped response box (Current Designs 4-Button Diamond Fiber Optic Response Pad) to make choices during the social task. Finally, a T1-weighted (T1w) MPRAGE sequence was acquired after the functional runs (192 slices, TR = 2000 ms, TE = 2.52 ms, flip angle = 12°, FOV = 256 mm, 1 mm isotropic voxels).

### Data Analysis

#### Image preprocessing

Preprocessing was performed using fMRIPrep 1.1.8 (Esteban et al., 2019), which is based on Nipype 1.1.3 (Gorgolewski et al., 2011) and Nilearn 0.4.2 (Abraham et al., 2014). Internal operations of fMRIPrep also use ANTs 2.1.0 (Avants et al., 2008), AFNI 17.2.07 (Cox, 1996), FSL 5.0.10 (Smith et al., 2004), and FreeSurfer 6.0.0 (Dale et al., 1999). The descriptions of anatomical and functional data preprocessing provided in the following two sections are based on the recommended citation boilerplate text that is generated by fMRIPrep and released under a CC0 license, with the intention that researchers reuse the text to facilitate clear and consistent descriptions of preprocessing steps, thereby enhancing the reproducibility of studies.

#### Anatomical data preprocessing

The T1w image was corrected for intensity non-uniformity using N4BiasFieldCorrection in ANTs 2.1.0, and used as T1w-reference throughout the workflow. The T1w-reference was then skull-stripped using antsBrainExtraction.sh (ANTs 2.1.0), using OASIS as target template. Brain surfaces were reconstructed using recon-all from FreeSurfer 6.0.0, and the brain mask estimated previously was refined with a custom variation of the method to reconcile ANTs-derived and FreeSurfer-derived segmentations of the cortical gray-matter of Mindboggle (Klein et al., 2017). Spatial normalization to the ICBM 152 Nonlinear Asymmetrical template version 2009c (MNI152NLin2009cAsym) was performed through nonlinear registration with antsRegistration (ANTs 2.1.0), using brain-extracted versions of both T1w volume and template. Brain tissue segmentation of cerebrospinal fluid (CSF), white-matter (WM) and gray-matter (GM) was performed on the brain-extracted T1w using fast (FSL 5.0.10).

#### Brain parcellation

Automated brain parcellation methods were used to identify our key regions of interest (right and left SPL) and to delineate other regions to be used in exploratory analyses. As mentioned in the preceding section, cortical surfaces for each participant were reconstructed by applying FreeSurfer’s anatomical parcellation algorithm to each participant’s T1w image. This process entails removal of non-brain tissue, automated segmentation of the cerebral cortex, subcortical white matter, brainstem, cerebellum, and deep gray matter structures, creating of a model of the participant’s cortical surface, and automatically parcellating this cortical surface model into discrete regions based on the participant’s cortical folding patterns. The Desikan-Killiany-Tourville (DKT) atlas (Klein and Tourville, 2012), implemented in FreeSurfer 6.0.0, was used to assign labels to cortical brain regions in each participant’s native space. This gyral-based atlas largely uses sulci to define the boundaries of adjacent gyri; as such, a particular gyral label in this atlas corresponds to both the gyrus itself and the adjacent banks of its limiting sulci. Using this procedure, each participant’s cerebral cortex was parcellated into 31 regions in each hemisphere. Six subcortical gray matter structures in each hemisphere, delineated using the tissue segmentation procedure described above were also retained for exploratory analyses (i.e., hippocampus, amygdala, caudate nucleus, palladium, putamen, nucleus accumbens). Thus, while our primary analyses concerned the right and left SPL, exploratory analyses spanned 74 anatomically defined brain regions.

#### Functional data preprocessing

For each of the 9 blood oxygen level-dependent (BOLD) runs per subject (across all tasks), the following preprocessing was performed. First, a reference volume and its skull-stripped version were generated using a custom methodology of fMRIPrep. A deformation field to correct for susceptibility distortions was estimated based on two echo-planar imaging (EPI) references with opposing phase-encoding directions, using 3dQwarp (AFNI 17.2.07). Based on the estimated susceptibility distortion, an unwarped BOLD reference was calculated for a more accurate co-registration with the anatomical reference. The BOLD reference was then co-registered to the T1w reference using bbregister (FreeSurfer 6.0.0), which implements boundary-based registration. Co-registration was configured with nine degrees of freedom to account for distortions remaining in the BOLD reference. Head-motion parameters with respect to the BOLD reference (transformation matrices, and six corresponding rotation and translation parameters) were estimated before any spatiotemporal filtering using mcflirt (FSL 5.0.10). The BOLD time-series were resampled onto their original, native space by applying a single, composite transform to correct for head-motion and susceptibility distortions. These resampled BOLD time-series will be referred to as preprocessed BOLD. Automatic removal of motion artifacts using independent component analysis (ICA-AROMA) was performed on the preprocessed BOLD in MNI space time-series after spatial smoothing with an isotropic, Gaussian kernel of 6mm FWHM (full-width half-maximum). Several confounding time-series were calculated based on the preprocessed BOLD, including framewise displacement (FD), and three region-wise global signals extracted within the CSF, the WM, and the whole-brain masks. FD was calculated for each functional run using implementations in Nipype. Additionally, a set of physiological regressors were extracted to allow for component-based noise correction (CompCor). Principal components were estimated after high-pass filtering the preprocessed BOLD time-series (using a discrete cosine filter with 128s cut-off) for anatomical CompCor (aCompCor). A subcortical mask was obtained by heavily eroding the brain mask, which ensured it does not include cortical GM regions. Six aCompCor components were calculated within the intersection of the aforementioned mask and the union of CSF and WM masks calculated in T1w space, after their projection to the native space of each functional run (using the inverse BOLD-to-T1w transformation). In addition, non-steady volumes identified by fMRIPrep were removed from data prior to subsequent analyses.

#### First-level analysis

We conducted run-wise analyses by fitting a general linear model (GLM) to each run of the fMRI data using Nipype (Gorgolewski et al., 2011) to estimate the BOLD response evoked for each direction of attentional shift in social knowledge (Task II) and external space (Task III). The following confounding variables (estimated during the preprocessing steps described in the preceding section) were included in the model as nuisance regressors: three translational motion parameters, three rotational motion parameters, three global signals extracted within the CSF, WM, and whole-brain masks, the set of aCompCor regressors, the set of discrete cosine basis functions that were used when extracting the aCompCor regressors, and the motion-related components identified by ICA-AROMA. All regressors of interest were convolved with a double gamma hemodynamic response function (HRF); the temporal derivative of the HRF was also included in the model. The *t*-statistic maps (i.e., maps of beta coefficients divided by their standard error estimates) resulting from these run-wise analyses were used for pattern similarity analysis (our primary analyses), as described in more detail below. In addition, we conducted trial-wise analyses prior to conducting classification analyses, in order to provide more training data to the SVM algorithm. Specifically, we repeated the above-described procedure modeling the response for each trial (rather than for each condition/direction of attentional shift) separately.

#### Pattern similarity analysis

We conducted a pattern similarity analysis in the SPL as our primary analysis. For each of Task II (the social task) and Task III (the spatial task), we calculated and compared the similarity of the neural response patterns (*Sim*) evoked when shifting attention in the same (matching) direction (*Sim_matching_*) and in opposite (mismatching) directions (*Sim_mismatching_*) across runs. *Sim_matching_* was calculated as the normalized mean correlation between patterns evoked during attentional shifts in the same direction across different runs (for example, between patterns evoked during “upward” shifts of attention in social knowledge in run 1 and “upward” shifts of attention in social knowledge in run 2). Contrastingly, *Sim_mismatching_* was calculated as the normalized mean correlation between patterns evoked during attentional shifts in opposing directions across different runs (for example, between patterns evoked during “upward” shifts of attention in social knowledge in run 1 and “downward” shifts of attention in social knowledge in run 2). More specifically, to calculate *Sim* scores, correlation coefficients were computed between the two types of patterns (matched and mismatched directions of attentional shifts) across each pair of runs, then transformed using Fisher’s *z* transformation, then normalized (i.e., *z*-scored) within participant, and finally, averaged across cases where attentional shift directions were matched and across cases where they were mismatched, within participant. If response patterns in a given brain region encode the direction of attentional shifts in social knowledge, then response patterns evoked during shifts of attention in the same direction should be more similar than those evoked during shifts of attention in opposing directions (i.e., *Sim_matching_* should be greater than *Sim_mismatching_*). Thus, we compared *Sim_matching_* to *Sim_mismatching_* through a paired samples *t*-test.

When conducting pattern similarity analyses across (rather than within) the social and spatial tasks, *Sim_matching_* was calculated as the mean normalized correlation between response patterns evoked during attentional shifts in matching directions across runs from different tasks (for example, between “upward” shifts of attention in social knowledge in run 1 of the social task, and upward shifts of visual attention in run 7 of the spatial task). *Sim_mismatching_* was calculated as the mean normalized correlation between response patterns evoked during attentional shifts in mismatching directions across runs from different tasks (for example, between patterns evoked during “upward” shifts of attention in social knowledge in run 1 of the social task, and downward shifts of visual attention in run 7 of the spatial task). As in the within-task pattern similarity analyses described in the preceding paragraph, correlation coefficients were again transformed using Fisher’s *z*-transformation, *z*-scored within participant, then averaged within conditions and participants; *Sim_matching_* and *Sim_mismatching_* were compared using a paired samples *t*-test.

After conducting our primary analyses in the left and right SPL, we repeated these pattern similarity analyses within each region of the DKT cortical parcellation, and in subcortical grey matter structures, using FDR-correction to account for multiple comparisons across regions. Additional exploratory whole-brain analyses were performed with searchlights, and further Bayesian hypothesis testing was also performed for cross-task analyses based on parcellations (described in the subsequent section).

#### Bayesian hypothesis testing

Following the paired samples *t*-tests that compared *Sim_matching_* and *Sim_mismatching_* in cross-domain pattern similarity analyses, we found that although certain brain regions encoded the “directions” of attentional shifts in both social knowledge and space (see Fig. 7a and Fig. 7b), cross-domain analyses did not show that they encoded both types of information in the same way (i.e., *Sim_matching_* and *Sim_mismatching_* were not significantly different in cross-domain pattern similarity analyses). Therefore, additional Bayesian hypothesis tests were conducted to assess the relative fitness of the null model (*Sim_matching_* = *Sim_mismatching_*) versus a two-sided alternative model (*Sim_matching_* ≠ *Sim_mismatching_*) in such regions. A Bayesian factor BF_10_ was calculated for each of those regions with the R package BayesFactor (Morey and Rouder, 2015). According to the current standards for Bayesian analyses (e.g., Gronau et al., 2020; van Doorn et al., 2019), we chose a Cauchy distribution with a spread 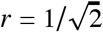 as the prior.

#### Searchlight analyses

Whole-brain searchlights were run on the social task and the spatial task, both separately and across the two tasks. The searchlights were run in each subject’s native space, within a dilated cortical ribbon mask. The cortical ribbon masks for each subject were obtained with FreeSurfer 6.0.0 based on their structural scans, downsampled to the same resolution as the functional scans, and then dilated with a spherical kernel of 3 mm. The searchlights were implemented in PyMVPA (Hanke et al., 2009) as spheres with radii of 4 voxels. Both the searchlight-based pattern similarity analyses (described in the preceding section) and additional exploratory classification analyses (described in the subsequent section) were run within each searchlight sphere. Maps of results from these analyses were smoothed with a gaussian kernel of 3 mm FWHM and transformed to MNI space prior to group analyses. The results of searchlight analyses were tested against either 0 (for pattern similarity analyses) or 0.5 (chance accuracy for classifications). FSL “randomise” was used to perform permutation tests and Threshold-Free Cluster Enhancement (TFCE) was used for multiple comparisons correction.

#### Classification analyses

Searchlight-based classifications were implemented in PyMVPA. Within each searchlight sphere, a linear support vector machine (SVM) classifier was trained on the social task (Task II) to classify the directions of attentional shifts (“upward” or “downward”), and its classification accuracy was tested, without further training, on the corresponding directions in the spatial task (Task III); another SVM classifier was trained on the spatial task and tested on the social task. Here, the correspondence between the attentional shift directions across the two tasks was determined individually, based on how each participant reconstructed the social hierarchy (see Fig. 4 and Supplementary file 5). All SVM classifiers, except in exploratory supplementary analyses described in Supplementary file 2, were trained with the default hyperparameters as implemented in the *LinearCSVMC* class of PyMVPA. In additional exploratory analyses (Supplementary file 2), we examined results using various classification algorithms and using a grid search procedure for hyperparameter-tuning (results of these exploratory analyses were consistent with those of the cross-task classification analyses described in the main text).

To better understand the results of this classification analysis, we conducted a separate analysis of within-task decoding (see Supplementary file 1).

#### Visualization

Visualizations of neuroimaging data throughout this paper were implemented with PySurfer (Waskom et al., 2020) and FSL (Smith et al., 2004).

## Footnotes

1 For participants who responded with diagonal representations, we related their multivoxel response patterns during social hierarchy navigation to *vertical* shifts of spatial attention in the main analyses. We conducted additional analyses where the response patterns were related to *horizontal* shifts of spatial attention, and this method yielded largely convergent results to those of our main analyses (see supplementary file 5).

2 We separately analyzed how the amount of “distance traveled” in social hierarchy knowledge (i.e., the magnitudes of attentional shifts) is reflected in multivoxel pattern similarity (see Supplementary file 6).

