## Supplementary file 1. Within-domain decoding for "How does the brain navigate knowledge of social relations? Testing for shared neural mechanisms for shifting attention in space and social knowledge"

As described in the main text, cross-domain decoding analyses identified no regions that encoded shifts of attention in space and social knowledge similarly using either a searchlight or a parcel-based approach, mirroring the results of the cross-domain pattern similarity analyses. Unsuccessful cross-domain decoding could reflect either attentional shift directions being encoded in different ways across domains, or our classification method having insufficient power to detect systematic distinctions between shifts of attention in different directions within domains. In other words, chance-level cross-domain decoding could stem from classifiers being unable to learn reliable distinctions within domains, or from classifiers learning reliable distinctions within domains that do not generalize across domains. To arbitrate between these possibilities, we conducted a within-domain decoding analysis.

**Data analysis.** This within-domain analysis was implemented similarly to the cross-domain decoding analyses described in the main text, with whole-brain searchlights and linear support vector machine (SVM) classifiers (see Methods for more details). Instead of training classifiers on either the social task or the spatial task and then testing classifiers on the left-out task, this analysis was carried out on each of the two tasks separately. For each task, specifically, we used leave-one-run-out cross-validation within each searchlight sphere, such that classifiers were trained on data from all fMRI runs except one, and then tested for their performance on the left-out run. Six-fold cross-validation was used for the social task (6 runs) and 2-fold cross-validation was used for the spatial task (2 runs), so that the data from each run was left out exactly once; classifier accuracies were then averaged across folds.

**Results.** Results from this within-task decoding analysis showed significantly above-chance accuracy for both the social task and the spatial task (Fig. S1a), in the superior parietal cortex, as well as in regions of the occipital cortex.

These results resembled the results we obtained through pattern similarity analyses (Fig. 6b/Fig. S1b). This suggests that this classification method was able to detect information signaled in multivoxel patterns about the direction of spatial and social attentional shifts in many of the same brain regions identified using the methods employed in our main analysis. This analysis generally resulted in smaller significant areas compared to the pattern similarity analysis, which may be due to the relatively small amount of training data available to the machine learning classifiers.

Using the same classification method, however, the classifiers were unable to achieve above-chance  in the cross-domain decoding analysis, as described in the main text. This indicates that the lack of cross-domain decoding success in the SPL was unlikely due to an inability of the current statistical methods to detect within-domain distinctions. Instead, SPL may not encode attentional shift directions in social and the spatial contents in the same way.

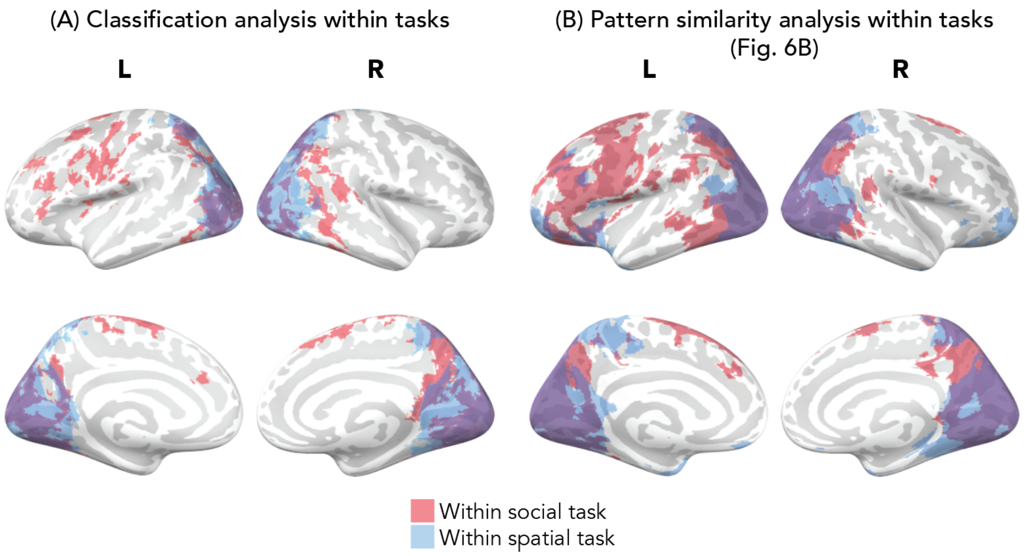


**Figure S1. Within-task classification analyses yielded similar results to within-task pattern similarity analyses.** We conducted a decoding analysis within each of the social and spatial domains in order to examine if the lack of significant results in the cross-domain decoding analysis (particularly in the superior parietal lobule) was due to an inability of classifiers to learn distinctions within tasks, or to within-task decision boundaries not generalizing across tasks. The results from this within-domain analysis (**A**) showed significantly above-chance accuracy in brain areas in much of the occipital and parietal lobule, largely replicating the results from the within-domain pattern similarity analysis (**B**). Panel B shows the results from the pattern similarity analysis, i.e., Fig. 6b, to provide a side-by-side comparison. All visualized results are significant at a corrected threshold of $p < .01$ (one-tailed).
