## Supplementary file 2. Hyperparameter tuning for cross-domain classification analysis for "How does the brain navigate knowledge of social relations? Testing for shared neural mechanisms for shifting attention in space and social knowledge"

For the classification analyses described in both the main text and Supplementary file 1, we used the default implementation and hyperparameters of linear SVM classifiers in PyMVPA (i.e., the *LinearCSVMC* class with *C* = -1, such that values are automatically scaled to the norm of the data). As the cross-domain classification analyses did not result in any significant result, we conducted additional exploratory analyses to examine if the same results would be achieved with hyperparameter tuning for the SVM classifiers, or with other classifiers. The following analyses were performed with Nilearn within each region of DKT cortical parcellation.

For SVM classifiers, we tried both linear and radial basis function (RBF) kernels, and selected hyperparameters for each with a grid search approach. The hyperparameters ranged as follows: *C*: 0.01, 0.1, 1, 10, 100; *gamma* (for RBF kernel only): 0.0001, 0.001, 0.01, 0.1, 1, 10, 100. The best hyperparameters were selected based on the results of 5-fold cross validations within either the social or spatial tasks, and then the best performing model from one task was trained on all of the data from that task, tested on data from the other task. For example, to select the best *C* for the linear SVM model, in one of the iterations, we built a linear SVM model using one of the possible *C* values, trained it on 80% of the data of the social task, and validated the model performance on the other 20% of the social task data. The performance for this *C* value is then calculated as the mean accuracy after 5-fold cross validation within the social task data. This procedure is then repeated for each possible *C* value, and the *C* value that resulted in the highest classification accuracy within the social task data was selected. The model with the best-performing parameters within the social task data was then trained on all data from the social task and tested on the data from the spatial task. Similar procedures were repeated for the SVM with RBF kernel, except that all combinations of *C* and *gamma* were iterated through in order to find the best model.

Besides SVM classifiers, we also further tested the performance of logistic regression. The logistic regression model was trained with stochastic gradient descent learning, and with L1 regularization (absolute norm), or L2 regularization (squared euclidean norm), or a combination of both (elastic net). Here, we integrated both the regularization method (L1, L2, or elastic net) and the strength of regularization as hyperparameters in grid search. The strength of regularization was selected among $10^{-6}$, $10^{-5}$, $10^{-4}$, 0.001, 0.01, and 0.1. That is, the optimal regularization method and the strength of regularization were determined based on the best-performing model from grid search results, in the same manner as *C* and *gamma* in the SVM classifier.

In additional exploratory analyses, we added feature selection as the first step in the pipeline to remove relatively non-informative features (i.e., voxels) before the SVM classification and logistic regression. We separately used either a univariate feature selection approach or a principal component analysis (PCA) approach for this purpose. For the univariate feature selection, ANOVA F-values were computed between each voxel and the class labels, and the voxels with the top *K*% scores were selected for later model fitting. Here, the percentage *K* was determined as a hyperparameter in grid search, which was selected among 25%, 40%, 55%, 70%, 85%, and 100% (no feature selection is done when *K* = 100%). Separately, with the PCA approach, the number of principal components *N* was also integrated as a hyper-parameter in grid search. *N* was set such that the amount of variance that is explained by the principal components is just above 70%, 80%, 90%, or 95% -- again, the best number was determined based on the grid search results.

For each of these analyses, FDR correction was used to account for multiple comparisons across regions.

**Results.** None of the exploratory classification methods above resulted in a significantly above-chance cross-domain classification in any brain region. Taken together with the significant results from within-domain decoding using less powerful methods (i.e., SVM classifiers without hyperparameter tuning or feature selection; see Supplementary file 1), as well as the cross-domain pattern similarity, decoding, and Bayesian hypothesis testing results described in the main text, these null results suggest that it is unlikely that the “directions” of attentional shifts across the social knowledge and space are encoded in the same way by the brain.
