## Supplementary file 3. Eye-tracking study for "How does the brain navigate knowledge of social relations? Testing for shared neural mechanisms for shifting attention in space and social knowledge"

To confirm that the results observed in the social hierarchy navigation task did not merely result from distinctive eye movements during mental navigation of social knowledge, we ran a separate study with the same tasks, while participants' eye movements were tracked during the social task.

In this eye-tracking study, participants first completed the trial-and-error learning task, followed by 6 blocks of the social task if they passed an accuracy criterion in the learning task (i.e., no more than 3 errors in adjacent trials of at least 3 consecutive blocks, and no more than 2 errors in non-adjacent trials of at least 3 consecutive blocks). We only analyzed data from participants who had satisfactory accuracy in the social task (>70%). Both paradigms are exactly the same as in the behavioral portion of the fMRI study.

**Participants.** Sixty-four participants were recruited from the UCLA community; 46 people passed the accuracy criterion in the learning task and thus continued to the social task (i.e., the eye-tracking portion), but 4 people were unable to complete the social task due to equipment malfunction. Out of the 42 people who completed the study, 12 were excluded from data analysis due to unsatisfactory accuracy (11 people) or being non-native English speakers (1 person; only native English speakers were included in the eye-tracking study to mirror the selection criteria for the fMRI study as closely as possible). The remaining 30 participants (10 males, 20 females) were native English speakers between the ages of 19 and 28 ($M = 20.5$)[1].

**Data acquisition.** Eye movements were recorded using an EyeTribe tracker at a sampling rate of 60 Hz. Participants were seated with their chin on a chinrest, which was placed approximately 50 cm from the screen. The social task was divided into two parts of 3 blocks each; participants had the chance to take a break in between the first and second part of the task. The eye tracker was calibrated at the beginning of each part using the EyeTribe UI and 16-point grid.

**Data analysis.** We extracted the raw coordinates of eye gaze positions during the 6000-ms mental navigation period of each trial, and calculated the median $x$ and $y$ coordinates for each trial. As described in the Methods section, the social task had two types of trials, where participants either navigated "up" (to find a more powerful person) or "down" (to find a less powerful person) in their knowledge of the learned social hierarchy. The median coordinates of eye gaze positions were then averaged across the "up" and "down" mental navigation trials for each participant. A paired $t$-test was used to test for differences between eye gaze coordinates in trials involving mental navigation of social knowledge in different "directions."

**Results.** On the $x$-axis, the mean difference of gaze positions between trials involving mental navigation in different "directions" (i.e., navigating "up" vs. "down" in social knowledge) was 1.97 pixels ($SD = 11.62$, $t(29) = 0.93$, $p = 0.36$). On the $y$-axis, the mean difference was -1.95 pixels ($SD = 9.06$, $t(29) = -1.18$, $p = 0.25$). These results suggest no significant systematic

[1] The same pattern of results that is reported here was observed without any participant exclusions (on the x-axis, mean difference = 0.62 pixels, $SD = 15.66$, $t(41) = 0.26$, $p = 0.80$; on the y-axis, mean difference = -1.45 pixels, $SD = 10.74$, $t(41) = -0.88$, $p = 0.39$). However, to provide a more stringent test of systematic associations between eye movements and mental navigation of social knowledge in our fMRI task, we focus on the results obtained from a sample with characteristics that mirror our fMRI sample (i.e., native English speakers who performed the social task with high accuracy).

differences in eye movements between trials corresponding to different navigation directions in the social task.
