## Supplementary file 4. Reference face analysis for "How does the brain navigate knowledge of social relations? Testing for shared neural mechanisms for shifting attention in space and social knowledge"

We recognized that in the social hierarchy navigation task, certain faces in the hierarchy were more likely to appear as reference faces in trials of a particular "direction". For example, faces that are relatively powerful in the hierarchy are more likely to be reference faces in a trial where participants were asked to find a less powerful person, rather than a more powerful one. To account for this unequal representation of reference faces across conditions, we undersampled the data so that each face appeared for the same number of times, and then conducted the same pattern similarity analysis reported in the main text.

**Data analysis.** We kept 18 out of the 24 trials in each run of the social task in this analysis, where participants were given 1) the third and the seventh face in the hierarchy; in these trials, participants were asked to find another face that was 2 steps away in either direction, 2) the fourth and the sixth face; participants were asked to find another face that was 2 or 3 steps away in either direction, and 3) the fifth face; participants were asked to find another face that was 2, 3, or 4 steps away in either direction. Thus, in this undersampled dataset, each reference face appears the same number of times in trials corresponding to attention shifts in each "direction" in the social hierarchy.

**Results.** The results from this analysis (yellow areas in Fig. S2) are similar to the results based on analysis of all 24 trials (Fig. 6B; red areas in Fig. S2). Therefore, we concluded that the unequal representation of reference faces across conditions had limited impact on the results.

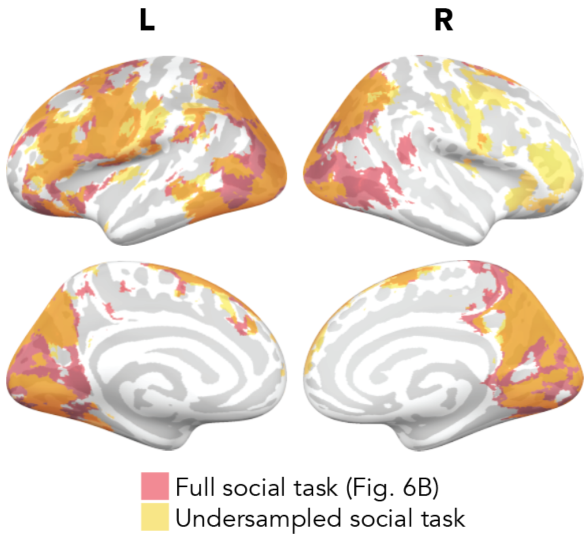


**Figure S2. Comparison between the results of full analysis (red) versus undersampled analysis (yellow) for the social hierarchy navigation task.** We undersampled the neuroimaging data for the social task in order to examine how much the unequal representation of reference faces, which cued the starting point of mental navigations in the social task, affected the results of the pattern similarity analyses. Red and yellow indicate areas that encoded "directions" of mental navigation in social knowledge, using the complete and undersampled neuroimaging data, respectively (orange indicates overlap).
