## Supplementary file 5. Alternative analyses for participants with diagonal representations of the social hierarchy for "How does the brain navigate knowledge of social relations? Testing for shared neural mechanisms for shifting attention in space and social knowledge"

In the hierarchy reconstruction task, four participants responded with a diagonal (or nearly diagonal) representation of the social hierarchy (see Fig. 4b). In the main cross-task analyses, we related these participants' brain response patterns in the social task to the *vertical* shifts of attention in the spatial task. Here, we conducted additional analyses while relating these four participants' brain response patterns in the social task to the *horizontal* shifts of attention in the spatial task (left-to-right for two participants and right-to-left for the other two). These additional analyses include the parcel- and searchlight-based pattern similarity analyses (see Methods).

For both the parcel- and searchlight-based pattern similarity analyses, results remain the same as the main analyses, i.e., no areas were identified to encode the attentional shifts in the social and spatial tasks in the same way.
