## Supplementary file 6. Analysis of the magnitude of attentional shifts within social knowledge for "How does the brain navigate knowledge of social relations? Testing for shared neural mechanisms for shifting attention in space and social knowledge"

During the social task (Task II), participants were asked to find target faces that were 2, 3, or 4 steps away from the given reference face. In the main analyses, we showed that SPL as well as other parietal and occipital regions encoded the "directions" of shifts of attention in social knowledge (i.e., whether participants were shifting their attention towards a more or less powerful person). Here, we analyzed how the brain encoded the "magnitudes" of such shifts of attention, i.e., the number of steps that was required for participants to shift their attention in each trial of the social task. However, note that the magnitude here is inherently confounded by task difficulty -- the farther away a target face is from the reference face, the more difficult it is for a participant to reach the correct answer. This was confirmed verbally by the participants in the pilot version of the study. Therefore, this analysis was performed only for exploratory purposes and the results should be interpreted with caution.

**Data analysis.** We performed a searchlight-based representational similarity analysis (RSA) to compare the relative similarity of the number of steps required for attentional shifts in different trials of the social task with the relative similarity of multivoxel response patterns during the 6-second attentional shift portion of the trials. Similar to the pattern similarity analyses focused on directions of attentional shifts described in the main text, we first obtained run-wise *t*-statistic maps in the first-level analysis (see Methods) corresponding to voxel-wise responses evoked by attentional shifts spanning 2, 3, and 4 "steps" in knowledge of the social hierarchy. At each searchlight center, local multivoxel response patterns evoked by attentional shifts of each magnitude (2, 3, and 4 "steps") were extracted. The representational similarity between local neural response patterns and a model signalling the relative magnitude of attentional shifts was computed by taking the Spearman correlation between the pairwise correlations of these local neural response patterns and the pairwise similarities (i.e., inverted absolute differences) of the corresponding number of steps of attentional shifts. The result maps were Fisher *z*-transformed, smoothed, compared against zero, and TFCE-corrected (see more details in Methods, searchlight analyses).

**Results.** As shown in Fig. S3, this analysis suggested that diffuse areas across the brain encode the magnitudes of attentional shifts in social knowledge, particularly in areas of lateral frontal and parietal cortex. As mentioned previously, magnitudes of attentional shifts in social knowledge have an inherent positive correlation with task difficulty. As such, it is unsurprising that this analysis implicated areas in, for example, dorsolateral prefrontal cortex, that are typically involved in storing and operating on the contents of working memory (Smith and Jonides, 1999) and in meeting cognitive demands more generally (Duncan and Owen, 2000). We note that the current study was not designed to dissociate task difficulty and the magnitudes of attentional shifts in social knowledge (as this study was focused on encoding *directions* of attentional shifts in knowledge of social relations); these results should be interpreted with caution and are presented to potentially inform future research.

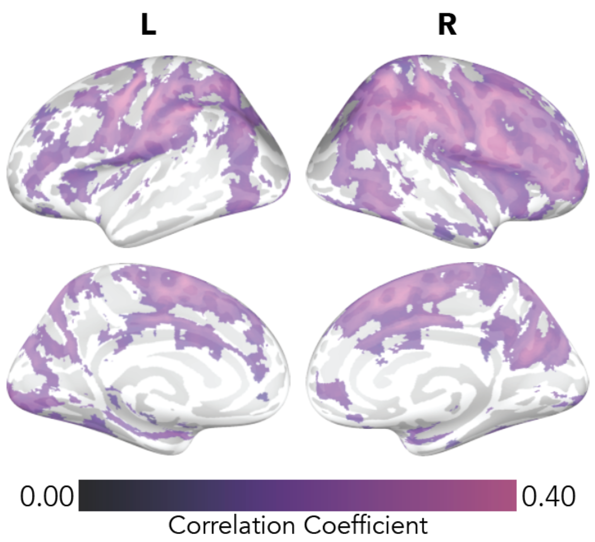


**Figure S3. Diffuse areas across the brain signaled the magnitude of attentional shifts within social knowledge (which could be due to task difficulty).** Purple-colored areas in this figure show where the relative similarity of local neural patterns is significantly related to the relative similarity of the magnitude of attentional shifts within social knowledge (i.e., where the correlation coefficients between similarity matrices based on the relative similarity of local neural patterns and of the magnitude of attentional shifts were significantly greater than 0, thresholded at $p < .001$, corrected for multiple comparisons with TFCE). These results should be interpreted with caution, given that the magnitude of attentional shifts is linked to overall task difficulty. We note that magnitudes of attentional shifts were balanced across attentional shift directions in all analyses reported in the main text.
