## Supplementary material for "How does the brain navigate knowledge of social relations? Testing for shared neural mechanisms for shifting attention in space and social knowledge": Figure 7 - figure supplement 1

| | Social | | | | | | Spatial | | | | | |
|---|---|---|---|---|---|---|---|---|---|---|---|---|
| | $\Delta Sim$ | *SD* | lower *CI* | $d_z$ | *t*(29) | FDR *p* | $\Delta Sim$ | *SD* | lower *CI* | $d_z$ | *t*(29) | FDR *p* |
| Pericalcarine cortex (L) | 0.074 | 0.089 | 0.046 | 0.83 | 4.54 | $4.75\times10^{-4}$ | 1.87 | 0.34 | 1.76 | 5.47 | 29.99 | $4.12\times10^{-22}$ |
| Cuneus cortex (R) | 0.097 | 0.091 | 0.069 | 1.07 | 5.86 | $4.28\times10^{-5}$ | 1.63 | 0.41 | 1.51 | 4.02 | 22.02 | $8.86\times10^{-19}$ |
| Lateral occipital cortex (L) | 0.091 | 0.080 | 0.066 | 1.13 | 6.21 | $3.30\times10^{-5}$ | 1.59 | 0.44 | 1.46 | 3.65 | 20.00 | $7.55\times10^{-18}$ |
| Lingual gyrus (L) | 0.082 | 0.11 | 0.047 | 0.72 | 3.97 | 0.0011 | 1.76 | 0.32 | 1.66 | 5.54 | 30.34 | $4.12\times10^{-22}$ |
| Cuneus cortex (L) | 0.058 | 0.081 | 0.033 | 0.72 | 3.93 | 0.0011 | 1.68 | 0.33 | 1.58 | 5.11 | 27.98 | $1.91\times10^{-21}$ |
| Lateral occipital cortex (R) | 0.065 | 0.066 | 0.045 | 1.00 | 5.47 | $8.32\times10^{-5}$ | 1.56 | 0.43 | 1.43 | 3.65 | 20.02 | $7.55\times10^{-18}$ |
| Lingual gyrus (R) | 0.092 | 0.13 | 0.051 | 0.70 | 3.82 | 0.0012 | 1.76 | 0.40 | 1.63 | 4.37 | 23.94 | $1.11\times10^{-19}$ |
| Pericalcarine cortex (R) | 0.11 | 0.15 | 0.064 | 0.75 | 4.12 | $9.47\times10^{-4}$ | 1.89 | 0.48 | 1.75 | 3.95 | 21.64 | $1.18\times10^{-18}$ |
| Superior parietal lobule (L) | 0.056 | 0.057 | 0.038 | 0.98 | 5.37 | $8.32\times10^{-5}$ | 0.92 | 0.49 | 0.77 | 1.90 | 10.41 | $1.07\times10^{-10}$ |
| Fusiform gyrus (L) | 0.053 | 0.060 | 0.035 | 0.89 | 4.87 | $2.66\times10^{-4}$ | 0.87 | 0.53 | 0.70 | 1.64 | 8.99 | $2.37\times10^{-9}$ |
| Superior parietal lobule (R) | 0.10 | 0.13 | 0.062 | 0.79 | 4.30 | $6.70\times10^{-4}$ | 0.83 | 0.53 | 0.67 | 1.58 | 8.65 | $4.93\times10^{-9}$ |
| Fusiform gyrus (R) | 0.038 | 0.065 | 0.018 | 0.58 | 3.19 | 0.0047 | 0.91 | 0.52 | 0.75 | 1.76 | 9.64 | $5.53\times10^{-10}$ |
| Inferior parietal lobule (R) | 0.055 | 0.074 | 0.032 | 0.75 | 4.10 | $9.47\times10^{-4}$ | 0.48 | 0.46 | 0.34 | 1.05 | 5.73 | $9.59\times10^{-6}$ |
| Inferior parietal lobule (L) | 0.050 | 0.063 | 0.030 | 0.79 | 4.31 | $6.70\times10^{-4}$ | 0.43 | 0.47 | 0.29 | 0.92 | 5.02 | $6.35\times10^{-5}$ |
| Precuneus (R) | 0.080 | 0.097 | 0.051 | 0.83 | 4.56 | $4.75\times10^{-4}$ | 0.40 | 0.52 | 0.23 | 0.76 | 4.14 | $5.55\times10^{-4}$ |
| Precuneus (L) | 0.061 | 0.082 | 0.035 | 0.74 | 4.05 | 0.0010 | 0.39 | 0.47 | 0.24 | 0.83 | 4.54 | $2.10\times10^{-4}$ |
| Insula (L) | 0.053 | 0.076 | 0.030 | 0.70 | 3.83 | 0.0012 | 0.23 | 0.40 | 0.10 | 0.57 | 3.14 | 0.0062 |
| Lateral orbitofrontal gyrus (L) | 0.045 | 0.10 | 0.012 | 0.43 | 2.34 | 0.025 | 0.41 | 0.49 | 0.26 | 0.85 | 4.65 | $1.64\times10^{-4}$ |
| Rostral middle frontal gyrus (L) | 0.055 | 0.079 | 0.031 | 0.70 | 3.82 | 0.0012 | 0.18 | 0.41 | 0.051 | 0.43 | 2.38 | 0.029 |
| Superior temporal gyrus (R) | 0.042 | 0.093 | 0.013 | 0.45 | 2.45 | 0.023 | 0.18 | 0.28 | 0.099 | 0.67 | 3.68 | 0.0018 |
| Middle temporal gyrus (R) | 0.041 | 0.065 | 0.021 | 0.63 | 3.47 | 0.0026 | 0.18 | 0.38 | 0.060 | 0.47 | 2.57 | 0.022 |
| Inferior temporal gyrus (L) | 0.046 | 0.075 | 0.023 | 0.62 | 3.37 | 0.0032 | 0.23 | 0.48 | 0.075 | 0.47 | 2.55 | 0.022 |
| Isthmus of cingulate cortex (L) | 0.039 | 0.091 | 0.011 | 0.43 | 2.35 | 0.025 | 0.23 | 0.35 | 0.12 | 0.66 | 3.62 | 0.002 |
| Lateral orbitofrontal gyrus (R) | 0.049 | 0.11 | 0.015 | 0.45 | 2.48 | 0.023 | 0.32 | 0.53 | 0.16 | 0.60 | 3.29 | 0.0044 |
| Orbital part of inferior frontal gyrus (L) | 0.052 | 0.082 | 0.027 | 0.63 | 3.48 | 0.0026 | 0.22 | 0.54 | 0.053 | 0.41 | 2.24 | 0.036 |
| Caudal middle frontal gyrus (L) | 0.046 | 0.073 | 0.024 | 0.63 | 3.47 | 0.0026 | 0.17 | 0.43 | 0.038 | 0.40 | 2.18 | 0.039 |
| Caudal middle frontal gyrus (R) | 0.032 | 0.065 | 0.011 | 0.49 | 2.66 | 0.016 | 0.24 | 0.47 | 0.091 | 0.50 | 2.75 | 0.016 |
| Putamen (R) | 0.074 | 0.13 | 0.035 | 0.59 | 3.25 | 0.0042 | 0.15 | 0.38 | 0.029 | 0.39 | 2.12 | 0.043 |
| Precentral gyrus (L) | 0.074 | 0.10 | 0.042 | 0.72 | 3.93 | 0.0011 | 0.10 | 0.37 | -0.016 | 0.27 | 1.46 | 0.12 |
| Parahippocampal gyrus (R) | 0.020 | 0.088 | -0.0070 | 0.23 | 1.26 | 0.14 | 0.25 | 0.32 | 0.15 | 0.79 | 4.35 | $3.34\times10^{-4}$ |
| Paracentral lobule (L) | 0.043 | 0.11 | 0.010 | 0.40 | 2.19 | 0.032 | 0.15 | 0.32 | 0.046 | 0.45 | 2.47 | 0.024 |
| Caudate nucleus (R) | 0.036 | 0.086 | 0.0094 | 0.42 | 2.30 | 0.027 | 0.16 | 0.38 | 0.039 | 0.41 | 2.27 | 0.035 |
| Triangular part of inferior frontal gyrus (L) | 0.056 | 0.079 | 0.032 | 0.72 | 3.93 | 0.0011 | 0.12 | 0.51 | -0.039 | 0.23 | 1.29 | 0.14 |
| Opercular part of inferior frontal gyrus (R) | 0.042 | 0.099 | 0.012 | 0.43 | 2.34 | 0.025 | 0.12 | 0.39 | 0.0051 | 0.32 | 1.77 | 0.077 |
| Postcentral gyrus (R) | 0.091 | 0.27 | 0.0062 | 0.33 | 1.82 | 0.061 | 0.18 | 0.51 | 0.025 | 0.36 | 1.97 | 0.055 |
| Inferior temporal gyrus (R) | 0.011 | 0.062 | -0.0085 | 0.17 | 0.95 | 0.21 | 0.24 | 0.36 | 0.13 | 0.67 | 3.67 | 0.0018 |
| Isthmus of cingulate cortex (R) | 0.022 | 0.085 | -0.0046 | 0.26 | 1.40 | 0.12 | 0.17 | 0.37 | 0.052 | 0.45 | 2.47 | 0.024 |

| | Social | | | | | | Spatial | | | | | |
|---|---|---|---|---|---|---|---|---|---|---|---|---|
| | *ΔSim* | *SD* | lower *CI* | $d_z$ | *t*(29) | FDR *p* | *ΔSim* | *SD* | lower *CI* | $d_z$ | *t*(29) | FDR *p* |
| Medial orbitofrontal gyrus (L) | 0.034 | 0.138 | -0.0086 | 0.25 | 1.36 | 0.12 | 0.21 | 0.46 | 0.071 | 0.46 | 2.55 | 0.022 |
| Amygdala (L) | 0.038 | 0.14 | -0.0062 | 0.27 | 1.46 | 0.11 | 0.17 | 0.40 | 0.044 | 0.42 | 2.30 | 0.033 |
| Middle temporal gyrus (L) | 0.047 | 0.11 | 0.014 | 0.44 | 2.42 | 0.024 | 0.09 | 0.37 | -0.022 | 0.25 | 1.37 | 0.13 |
| Supramarginal gyrus (R) | 0.058 | 0.14 | 0.016 | 0.43 | 2.35 | 0.025 | 0.12 | 0.45 | -0.024 | 0.26 | 1.41 | 0.13 |
| Triangular part of inferior frontal gyrus (R) | 0.026 | 0.086 | $-3.69\times10^{-4}$ | 0.31 | 1.68 | 0.079 | 0.16 | 0.43 | 0.021 | 0.36 | 1.96 | 0.055 |
| Superior frontal gyrus (R) | 0.026 | 0.059 | 0.0081 | 0.45 | 2.45 | 0.023 | 0.12 | 0.51 | -0.041 | 0.23 | 1.26 | 0.15 |
| Orbital part of inferior frontal gyrus (R) | 0.035 | 0.092 | 0.0063 | 0.38 | 2.07 | 0.039 | 0.13 | 0.50 | -0.020 | 0.27 | 1.48 | 0.12 |
| Posterior cingulate cortex (R) | 0.045 | 0.11 | 0.011 | 0.41 | 2.25 | 0.029 | 0.085 | 0.35 | -0.024 | 0.24 | 1.33 | 0.14 |
| Superior temporal gyrus (L) | 0.044 | 0.12 | 0.0070 | 0.37 | 2.02 | 0.042 | 0.11 | 0.41 | -0.018 | 0.27 | 1.46 | 0.12 |
| Opercular part of inferior frontal gyrus (L) | 0.057 | 0.090 | 0.029 | 0.64 | 3.49 | 0.0026 | 0.049 | 0.34 | -0.056 | 0.15 | 0.80 | 0.25 |
| Transverse temporal gyrus (L) | 0.026 | 0.126 | -0.013 | 0.21 | 1.13 | 0.17 | 0.12 | 0.33 | 0.021 | 0.38 | 2.06 | 0.048 |
| Precentral gyrus (R) | 0.11 | 0.26 | 0.025 | 0.41 | 2.23 | 0.030 | 0.090 | 0.48 | -0.058 | 0.19 | 1.04 | 0.20 |
| Insula (R) | 0.031 | 0.12 | -0.0063 | 0.26 | 1.41 | 0.12 | 0.15 | 0.51 | -0.011 | 0.29 | 1.58 | 0.11 |
| Rostral middle frontal gyrus (R) | 0.025 | 0.087 | -0.0020 | 0.29 | 1.57 | 0.094 | 0.13 | 0.51 | -0.028 | 0.26 | 1.40 | 0.13 |
| Entorhinal cortex (L) | 0.020 | 0.138 | -0.023 | 0.15 | 0.80 | 0.25 | 0.21 | 0.43 | 0.081 | 0.50 | 2.73 | 0.016 |
| Parahippocampal gyrus (L) | 0.033 | 0.141 | -0.011 | 0.23 | 1.27 | 0.14 | 0.12 | 0.43 | -0.011 | 0.28 | 1.56 | 0.11 |
| Posterior cingulate cortex (L) | 0.028 | 0.065 | 0.0081 | 0.43 | 2.38 | 0.025 | 0.052 | 0.35 | -0.057 | 0.15 | 0.81 | 0.25 |
| Medial orbitofrontal gyrus (R) | 0.038 | 0.14 | -0.0073 | 0.26 | 1.42 | 0.12 | 0.088 | 0.36 | -0.024 | 0.24 | 1.34 | 0.14 |
| Postcentral gyrus (L) | 0.048 | 0.099 | 0.017 | 0.48 | 2.63 | 0.017 | 0.069 | 0.58 | -0.11 | 0.12 | 0.65 | 0.29 |
| Paracentral lobule (R) | 0.083 | 0.24 | 0.0075 | 0.34 | 1.87 | 0.057 | 0.049 | 0.33 | -0.053 | 0.15 | 0.82 | 0.25 |
| Putamen (L) | 0.026 | 0.10 | -0.0067 | 0.25 | 1.35 | 0.12 | 0.085 | 0.51 | -0.074 | 0.17 | 0.91 | 0.23 |
| Supramarginal gyrus (L) | 0.044 | 0.082 | 0.019 | 0.54 | 2.96 | 0.0080 | -0.031 | 0.45 | -0.17 | -0.070 | -0.38 | 0.38 |
| Pallidum (L) | 0.051 | 0.13 | 0.010 | 0.39 | 2.11 | 0.036 | 0.041 | 0.46 | -0.10 | 0.089 | 0.49 | 0.34 |
| Hippocampus (R) | 0.0070 | 0.096 | -0.023 | 0.073 | 0.40 | 0.38 | 0.15 | 0.43 | 0.019 | 0.35 | 1.94 | 0.056 |
| Caudate nucleus (L) | -0.011 | 0.093 | -0.040 | -0.12 | -0.67 | 0.29 | 0.071 | 0.45 | -0.067 | 0.16 | 0.87 | 0.24 |
| Rostral anterior cingulate cortex (R) | 0.0094 | 0.13 | -0.032 | 0.071 | 0.39 | 0.38 | 0.12 | 0.51 | -0.041 | 0.23 | 1.25 | 0.15 |
| Caudal anterior cingulate cortex (L) | 0.013 | 0.073 | -0.0094 | 0.18 | 0.99 | 0.20 | -0.024 | 0.27 | -0.11 | -0.089 | -0.49 | 0.34 |
| Hippocampus (L) | -0.0016 | 0.075 | -0.025 | -0.021 | -0.11 | 0.47 | 0.18 | 0.44 | 0.042 | 0.41 | 2.23 | 0.036 |
| Amygdala (R) | 0.019 | 0.099 | -0.011 | 0.19 | 1.07 | 0.18 | -0.020 | 0.48 | -0.17 | -0.042 | -0.23 | 0.43 |
| Caudal anterior cingulate cortex (R) | -0.0025 | 0.093 | -0.031 | -0.026 | -0.14 | 0.46 | 0.12 | 0.43 | -0.015 | 0.27 | 1.51 | 0.12 |
| Superior frontal gyrus (L) | 0.038 | 0.049 | 0.023 | 0.78 | 4.29 | $6.70\times10^{-4}$ | 0.0037 | 0.44 | -0.13 | 0.0084 | 0.05 | 0.49 |
| Transverse temporal gyrus (R) | 0.012 | 0.21 | -0.054 | 0.056 | 0.30 | 0.41 | 0.061 | 0.55 | -0.11 | 0.11 | 0.60 | 0.31 |
| Nucleus accumbens (L) | 0.012 | 0.099 | -0.019 | 0.12 | 0.64 | 0.29 | -0.026 | 0.52 | -0.19 | -0.049 | -0.27 | 0.42 |
| Nucleus accumbens (R) | -0.0026 | 0.15 | -0.048 | -0.017 | -0.10 | 0.47 | 0.053 | 0.38 | -0.066 | 0.14 | 0.76 | 0.26 |
| Pallidum (R) | -0.0010 | 0.13 | -0.040 | -0.008 | -0.044 | 0.48 | 0.079 | 0.43 | -0.056 | 0.18 | 1.00 | 0.21 |
| Rostral anterior cingulate cortex (L) | 0.014 | 0.12 | -0.022 | 0.12 | 0.68 | 0.29 | 0.0047 | 0.52 | -0.16 | 0.0091 | 0.05 | 0.49 |
| Entorhinal cortex (R) | 0.0083 | 0.16 | -0.042 | 0.051 | 0.28 | 0.41 | $1.33\times10^{-4}$ | 0.49 | -0.15 | $2.74\times10^{-4}$ | 0.0015 | 0.50 |

**Figure 7 - figure supplement 1. Additional statistical results of parcel-based pattern-similarity analyses.** This figure is a more comprehensive version of Fig. 7. Both Fig. 7 and this figure contain results from within-domain pattern similarity analyses, which tested if the multivoxel response patterns for attentional shifts in matching directions were significantly more similar to each other compared to those in mismatching directions. Here, $\Delta Sim = Sim_{matching} - Sim_{mismatching}$, and $\Delta Sim$ was tested against zero. Fig. 7 only shows the brain regions that were significant at an FDR-corrected threshold of $p < 0.01$, one-tailed, in either the social task or the spatial task. This figure shows results for all brain regions, including those that did not surpass a one-tailed FDR-corrected threshold of $p < .01$ on one or both tasks (i.e., those not included in Fig. 7a).
